# PathGPS: Discover shared genetic architecture using biobank data

**DOI:** 10.1101/2022.05.01.490230

**Authors:** Zijun Gao, Trevor Hastie, Qingyuan Zhao

## Abstract

The increasing availability and scale of Genome Wide Association Studies (GWAS) bring new horizons for understanding biological mechanisms. PathGPS is an exploratory method that discovers genetic architecture using GWAS summary data. It can separate genetic components from unobserved environmental factors and extract clusters of genes and traits associated with the same biological pathways. When applying to a metabolomics dataset and the UK Biobank, PathGPS confirms several known gene-trait clusters and suggest many new hypotheses for future investigations.

---

Understanding the biological mechanisms by which genetic variation influences phenotypes is one of the primary challenges in human genetics^1^. Genome-wide association studies (GWAS) have successfully mapped thousands of genetic loci associated with complex human traits. However, it is extremely time-consuming and inefficient to investigate every identified association and validate the function^2^. Moreover, complex traits are usually highly polygenic and are associated with a large number of variants across the genome, each explaining only a small fraction of the genetic variance^3,4^. These difficulties have hindered the translation of GWAS findings into drug development and clinical applications^5^. When pooling data from multiple GWAS, recent studies found that many complex traits and diseases are associated with the same genomic loci^6^ and identified many pairs of traits with strong genetic correlation^7,8^. This indicates that many disease-causing variants may cluster into key pathways that drive several diseases at the same time^9^. Recent large-scale biobank data provide an exciting opportunity to discover shared genetic architecture between complex traits.

We start with a statistical model that assumes that most human traits are regulated by one or several genetic or environmental pathways (Figure 1a, Online Supplement 1). This model highlights several challenges to recover the shared genetic architecture. First, isolating the shared genetic contribution is difficult because the environmental factors are usually not observed. Second, it is often desirable to use data from multiple cohorts, but those cohorts may be sampled from different populations and measure different traits. Third, many traditional unsupervised learning methods are very sensitive to small perturbations to the data and tuning parameter selection.

**Figure 1:**
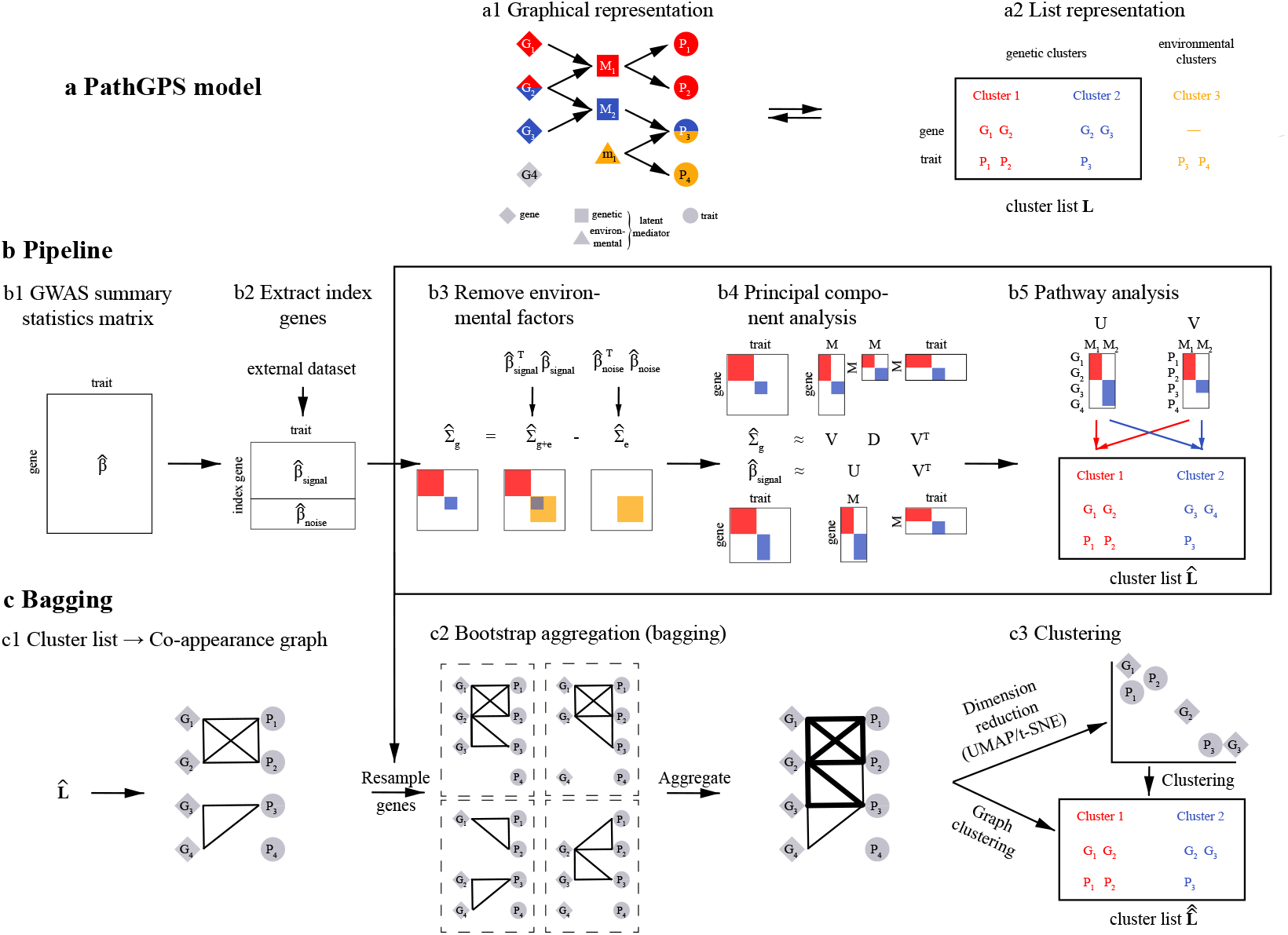
Illustrative summary of PathGPS. Panel **a** displays the graphical representation of our model for latent genetic and environmental pathways (**a1**) and the corresponding gene-trait clusters (**a2**). Panel **b** demonstrates how PathGPS takes a GWAS summary statistics matrix (**b1**), uses an external dataset to select signal and noise index SNPs (**b2**), subtracts the covariance matrix of the traits based on the noise index SNPs from that of the signal index SNPs to remove the environmental component (**b3**), applies principal component analysis to learn a low-rank strcuture (**b4**), and finally assigns SNPs and traits with non-zero loadings into clusters (**b5**). Panel **c** displays the proposed “bagging” procedure to stabilize the results: for each resample of the data, PathGPS first constructs a co-appearance graph by connecting SNPs and traits in the same gene-trait cluster (**c1**), then aggregates the co-appearance graphs by assigning edge weights proportional to the likelihood of vertices appearing in the same gene-trait cluster (**c2**), visualizes the aggregated graph and clusters in low-dimensional embeddings, and finally applies clustering algorithms to output the discovered biological pathways (**c3**).

We propose a new statistical method called PATHway discovery through Genome-Phenome Summary data (PathGPS) that can generate clusters of genes and traits associated with the same biological pathways (Figure 1b). By subtracting the empirical covariance matrix of traits computed using “noise” genes (genetic variants showing no or weak associations with the traits) from that using “signal” genes (genetic variants showing strong associations with some traits), PathGPS disentangles genetic mechanisms from environmental factors and overcomes the first challenge (Online Supplement 2.1, Corollary 1, Figure 2). PathGPS then applies principal component analysis (PCA) to the matrix difference and provides a low-rank representation of the genetic associations. We show that PathGPS is robust to batch effects (Online Supplement 2.1, Corollary 2) and thus overcomes the second challenge. Regarding the third challenge, PathGPS utilizes a novel bootstrap aggregating (“bagging”) method to stabilize the algorithm (Figure 1c, Online Supplement 2.2). By resampling the genes, PathGPS obtains a weighted graph that estimates the chance that a pair of variables (could be genes or traits) appear in the same pathway. Dimension reduction techniques and clustering algorithms are then applied to visualize this graph and produce the stabilized clusters.

**Figure 2:**
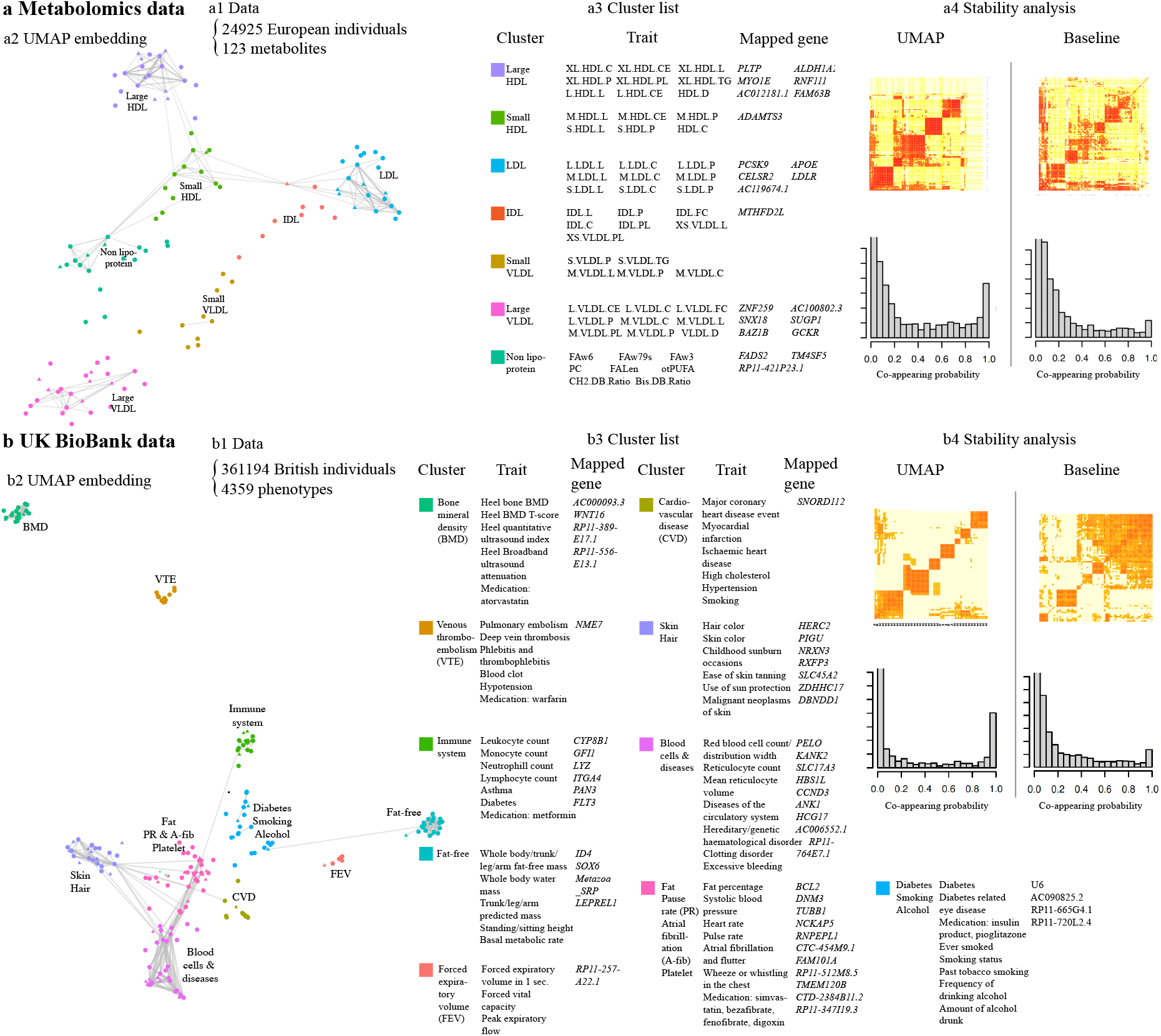
Applications of PathGPS. Panel **a** displays the summary of the metabolomics data (**a1**), the UMAP embeddings of 7 gene-trait clusters produced by PathGPS with co-appearance edge weights (**a2**), and representative traits and mapped genes in each cluster (**a3**). In (**a4**), we subsample traits without replacement, and PathGPS (UMAP) produces more consistent cluster memberships (Online Supplement 6.2) than the baseline method (Figure 1b5). Panel **b** displays the summary of the UK BioBank data (**b1**), the UMAP visualization (**b2**), and representative genes and traits of the 10 clusters produced by PathGPS (**b3**). PathGPS (UMAP) again produces more stable clusters (**b4**). The representative traits and mapped genes in (**a3**) and (**b3**) are selected manually (full list in Online Supplement 6 and Tables).

We evaluate the practical performance of PathGPS in a comprehensive simulation study with synthetic genes and traits generated from our statistical model. In the first set of simulations, we assess PathGPS’s ability in disentangling genetic and environmental influences (Online Supplement 3.1). We compare PathGPS with a baseline method that directly applies PCA to the empirical covariance matrix of traits using the signal genes. PathGPS successfully finds clusters of traits influenced by the same genetic pathways but not environmental factors, while the baseline method is unable to distinguish genetic effect from environmental effect. In the second set of simulations, we assess how bagging stabilizes the unsupervised learning algorithm. We compare PathGPS with (Figure 1c) and without bagging (Figure 1b). PathGPS with bagging produces significantly more robust results that is table across a range of reasonable hyper-parameters (Online Supplement 3.2, Figure 4).

We apply PathGPS to two real datasets: a genome-metabolome wide association study (*N* = 24925)^10^ and the UK BioBank (*N* = 361194)^11^ (Figure 2). We use *haploR*^12^ to map SNPs to genes. For the metabolomics dataset (Online Supplement 5.1), PathGPS produces 7 clusters which roughly correspond to large high-density lipoprotein (HDL), small HDL, low-density lipoprotein (LDL), intermediate-density lipoprotein (IDL), large very-low-density lipoprotein (VLDL), small VLDL, and non-lipoprotein measurements (Figure 2a). Thus, using genetic data only, PathGPS is able to recover the known taxonomy of circulating metabolites. Moreover, PathGPS confirms several known causal genes, such as *PLTP* as a regulator of HDL size^13^ and *PCSK9* as a regulator of LDL cholesterol^14^. PathGPS also proposes several biological hypotheses that are not as well established, including *RNF111* in relation to HDL^15^ and *TM4SF5* in relation to lipid measurements^16^. In fact, few gene-trait pairs suggested by PathGPS directly reach the genome-wide significant level after correcting for multiple testing, but the majority of the gene-trait pairs are at least moderately associated. This demonstrates the ability of PathGPS to associate a group of genes with a group of traits, when any single association is not strong enough.

For the UK BioBank, we investigate 175 traits that pass our quality control and are associated with at least one genetic variant (Online Supplement 5.2). PathGPS produces 10 clusters (Figure 2b3), among which 3 are closely related to some diseases (venous thromboembolism, cardiovascular diseases, and type 2 diabetes), and the other 7 contain biometric measurements, such as bone mineral density, immune system, fat-free mass, and skin or hair colors. In the UMAP visualization (Figure 2b2), the edges reflect high (top 350) co-appearances between vertex pairs and may offer insights into disease mechanisms. For instance, our analysis finds the medication simvastatin is closely related to high cholesterol and cardiovascular diseases (CVD), which is not surprising given that it is widely prescribed to reduce CVD risk^17^. We also find atorvastatin— another drug in the statin family—is highly associated with bone mineral density (BMD) and related traits. This finding is consistent with existing evidence that statin increases BMD^18,19^.

In addition, edges connecting monocyte, neutrophil, lymphocyte and diabetes, asthma diagnosis have high weights, suggesting connections between the immune system and the two common diseases. In particular, diabetes may be related to the immune system through multiple mechanisms: for example, hyperglycemia in diabetes may cause dysfunction of the immune response^20^. As for asthma, T lymphocytes are critical to the development of asthma^21^. The co-occurrence of diabetes and asthma may be attributed to the shared immunological pathways^22^. Regarding the genetic architecture, our analysis confirm many existing discoveries, such as the association between *HERC2* and hair color^23^, *PELO* and red blood cells^24^, and *NME7* and venous thromboembolism^25^. We also find less well established biological hypotheses, such as *BCL2* and Atrial fibrillation^26^, *GFI1* and lymphocyte cells^27^. The UMAP embedding provides further information beyond the cluster membership. For example, the cluster containing smoking, alcohol, and diabetes is adjacent to the cluster containing CVD, indicating a multifaceted health effect of alcohol consumption and tobacco usage.

In summary, PathGPS is a promising statistical tool to uncover clusters of genes and traits that correspond to latent biological pathways from multiomic and biobank data. When applied to the UK Biobank, PathGPS not only confirms many established genetic associations but also generates novel biological hypotheses. Finally, by grouping diseases with shared biological pathways, PathGPS can enhance our understanding of comorbidities and contribute to the development of comprehensive clinical practices.

## Methods

### Structural equation model of latent pathways

We describe the latent genetic and environmental pathways connecting genes (SNPs) and traits using linear structural equation models (SEMs). Consider SNPs ***X*** = (*X*_1_, …, *X*_*p*_) and traits ***Y*** = (*Y*_1_, …, *Y*_*q*_). We assume the traits are influenced by the SNPs through latent genetic mediators ***M*** = (*M*_1_, …, *M*_*r*_). Meanwhile, the traits are also affected by unobserved environmental mediators ***m*** = (*m*_1_, …, *m*_*s*_). Mathematically, we adopt the linear SEM (Figure 1a1 displays the graphical representation)

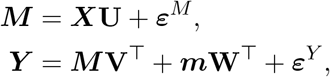

where **U** ∈ ℝ^*p×r*^, **V** *∈*R^*q×r*^, **W** ∈ ℝ^*q×s*^ are coefficient matrices, and ***ε***^*M*^ ∈ ℝ ^*r*^, ***ε***^*Y*^ ∈ ℝ ^*q*^ denote zero-mean errors in the genetic mediators ***M*** and traits ***Y***, respectively. Concerning that genetic pathways usually only include a small number of SNPs and traits, the coefficient matrices **U, V** are often sparse. We remark that the above SEM does not include the interaction of genetic and environmental pathways. Given that we only have access to summary statistics (details below), the SEMs with and without interaction are not identifiable and thus we cling to the simpler version without interaction.

### Genome-wide association studies summary statistics

Our analysis builds upon the genome-wide association studies (GWAS) summary statistics due to their wider availability compared to individual-level data *{*(***X, Y***)*}*. In particular, we focus on the effect estimates (marginal associations) 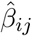 of SNP-trait pairs (*X*_*i*_, *Y*_*j*_) obtained from simple linear regressions. We select and work with approximately independent SNPs—index SNPs. As shown in the supplementary materials section 1, the marginal associations of index SNPs satisfy

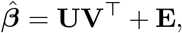

where **E** is a zero-mean matrix stemming from the environmental mediators ***m*** and the errors ***ε***^*M*^, ***ε***^*Y*^. Since both **U** and **V** are sparse and of rank *r* ≪ *p, q*, the marginal association matrix 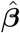 is centered around a sparse low-rank matrix. The low-rank and sparse structures are crucial to the development of the following statistical estimation procedures.

## Discovery of gene-trait clusters

Our goal is to discover genetic pathways that start from SNPs, pass through latent genetic mediators, and end in observed traits. Since the genetic mediators are unobserved, we look for clusters of SNPs and traits that are potentially related to the same underlying genetic mediator. For example, in Figure 1a1, there are two genetic pathways : *X*_1_, *X*_2_ → *M*_1_ → *Y*_1_, *Y*_2_ in red and *X*_2_, *X*_3_ → *M*_2_ → *Y*_3_ in blue. The corresponding gene-trait clusters are *{X*_1_, *X*_2_, *Y*_1_, *Y*_2_*}* and *{X*_2_, *X*_3_, *Y*_3_*}* (Figure 1a2). Under the notations of the proposed SEM, a desirable cluster should consist of the genes and traits with non-zero loadings in a matched column pair of **U** and **V**.

We reconstruct gene-trait clusters from the marginal association estimates 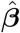. We first employ the low-rank structure of **U, V** to learn their low-dimensional column spaces. We next use the sparsity of **U, V** to find sparse matrices **Û**, 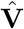 in the estimated column spaces. Finally, we assign genes and traits with non-zero loadings in a matched column pair of **Û**, 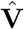 to a gene-trait cluster. Below we discuss the major estimation steps. to a gene-trait cluster.

### Column space recovery

We estimate the column spaces corresponding to the genetic pathways based on 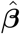 Note that 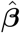 contains both genetic and environmental components (Online Supplement 2.1, Proposition 1).We use the marginal association estimates 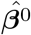 of noise SNPs—SNPs that have no statistically significant association with any traits—to estimate the non-genetic structure, and remove it from the marginal association estimates 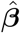 of signal SNPs—SNPs that are significant to at least one trait. The subtraction step removes the environmental factors and keeps the genetic factors (Online Supplement 2.1, Corollary 1).

In particular, we estimate the column space of **V** by the space spanned by the top *r* eigenvectors, denoted by 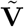, of

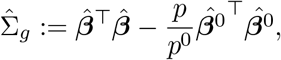

where *p, p*^0^ denote the number of signal and noise SNPs, respectively. Upon obtaining 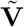, we estimate the column space of **U** by that of 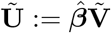. We remark that the number of eigen-vectors *r* is a hyper-parameter corresponding to the number of latent genetic mediators. In real data analyses, we plot the eigen-values of 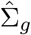 and let *r* be the visual “elbow-point”.

### Sparse coefficient matrix recovery

Based on the estimated column spaces of **U** and **V**, we propose methods to find **Û**, 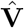 that are sparse and form the bases of the estimated spaces. In particular, we start from the **Û**, 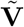 obtained in the column space recovery, and look for the transformation matrix **R** ∈ ℝ^*r×r*^ such that 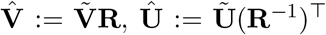 have sparse columns. There are a number of readily available methods from factor analysis to search for the appropriate transformation matrix **R**, and we use the varimax criterion^28^ in our analysis.

### Construction of co-appearance graph by SNP bootstrapping

Several issues may undermine the reliability of the discovery of gene-trait clusters: the evolving set of measured traits, the data pre-processing procedures (including the selection of signal and noise SNPs), and the choices of hyper-parameters (including the number of latent genetic mediators *r*). We posit a bootstrap aggregation (bagging) approach to stabilize the pipeline and make the results more replicable. The idea is to perturb the entire procedure many times and then aggregate the results using the co-appearance graph as described below.

### SNP bootstrapping

In a bootstrap trial, we resample the same number *p* of signal SNPs with replacement and obtain the bootstrap signal marginal association estimates 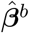. We extract a list ℒ_*b*_ of gene-trait clusters from 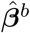 and 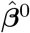 using the aforementioned approach. The marginal association estimates 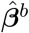 may have duplicate rows due to the sampling with replacement, and we will remove the duplicates in the co-appearance graph aggregation. We repeat the trial *B* times and get a collection of cluster lists (Figure 1c2 describes the bootstrap process). We remark that the SNP bootstrapping is only one way of perturbing the datasets, and other methods like sub-sampling SNPs without replacement also serve the goal.

### Co-appearance graph aggregation

We introduce the co-appearance frequency for a gene-trait pair, and the definition can be easily generalized to gene-gene pairs and trait-trait pairs. Let (*X*_*i*_, *Y*_*j*_) be a gene-trait pair, and we define the co-appearance frequency as

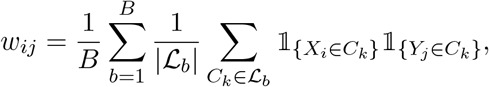

where *C*_*k*_ denotes the *k*-th cluster in the list ℒ_*b*_ and | ℒ_*b*_| stands for the number of clusters in the list. The co-appearance frequency describes how frequently a pair of gene and trait appear in the same cluster: if a pair of gene and trait always show up in the same cluster, the pair will have a high co-appearance frequency, and we are more confident that the gene and the trait lie in the same biological pathway.

With all the co-appearance frequencies, we can construct a weighted undirected graph of SNPs and traits to aggregate the cluster lists of bootstrap samples. We consider a graph where each node represents a SNP or a trait, and we weight each edge by the co-appearance frequency (Figure 1c3). The weighted graph is convenient for downstream analyses, including dimension reduction and clustering. Compared with the one-shot gene-trait cluster discovery, the bagging procedure underlying the co-appearance graph reduces several sources of instability (Online Supplement 3.2).

## Simulation study: stability analysis

In practice, datasets with moderately different collections of traits may address the same biological question, and it is desirable to arrive at consistent conclusions despite the differences in trait collection. We provide details of the stability analysis applied to the metabolomics data (Figure 2a4). The UK BioBank data is handled similarly (Figure 2b4). We start from the marginal association estimate matrix with a total of 105 traits. In each trial, we subsample 50 traits without replacement. We then compare the co-appearance weights obtained with (Figure 2a4 UMAP) and without (Figure 2a4 Baseline) bootstrap aggregation. If bagging increases the stability of the results, it is expected that the histograms of co-appearance weights will have sharper spikes around 0 and 1. Close-to-one co-appearance weights indicate the associated pairs always fall into the same cluster and close-to-zero values imply the associated pairs always end up in different clusters.

## Data availability

The main metabolomics dataset^10^ contains 123 metabolites and around 1.3 ×10^7^ SNPs. We use an external dataset^29^ of 72 metabolites and the PLINK software^30^ to select independent index SNPs. The detailed pre-processing procedures of the metabolomics data are summarized in Figure 5 of the supplementary materials.

The main UK BioBank data (UKBB GWAS Imputed v3) is generated by Neale Lab (http://www.nealelab.is/uk-biobank). The dataset is based on 3.6 *×* 10^5^ samples of white-British ancestry and 1.3*×*10^7^ QC-passing SNPs. The traits are a mixture of lab measurements, diagnoses and medications, and living habits. We use the summary statistics based on female subjects and the PLINK software to select index SNPs, and derive the clustering results from the summary statistics of the male subjects. The detailed pre-processing procedures are displayed in Figure 6 of the supplementary materials.

## Code availability

All methods described in this paper are implemented in and are available at https://github.com/ZijunGao/PathGPS.

## Acknowledgements

Qingyuan Zhao is supported by the Isaac Newton Trust and EPSRC grant EP/V049968/1. Zijun Gao and Trevor Hastie are partially supported by grants DMS 2013736 and IIS 1837931 from the (US) National Science Foundation and grant 5R01 EB 001988-21 from the (US) National Institutes of Health.

For the purpose of open access, the authors have applied a Creative Commons Attribution (CC BY) license to any Author Accepted Manuscript version arising from this submission.

## Notes

### Competing Interest Statement

The authors have declared no competing interest.

