## Supplementary material for "PathGPS: Discover shared genetic architecture using biobank data": Online Supplement

May 1, 2022

[*]Department of Statistics, Stanford University, CA, USA. Email: {zijungao}@stanford.edu.

[†]Department of Statistics and Department of Biomedical Data Science, Stanford University, CA, USA. Email: {hastie}@stanford.edu.

[‡]Statistical Laboratory, Department of Pure Mathematics and Mathematical Statistics, University of Cambridge, UK. Email: {qyzhao}@statslab.cam.ac.uk.

### 1 Problem formulation

Consider $p$ SNPs $\boldsymbol{X} = (X_1, \ldots, X_p)$ and $q$ traits $\boldsymbol{Y} = (Y_1, \ldots, Y_q)$. We assume the traits are influenced by the SNPs through $r$ latent genetic mediators $\boldsymbol{M} = (M_1, \ldots, M_r)$. Meanwhile, the traits are also affected by $s$ unobserved environmental mediators $\boldsymbol{m} = (m_1, \ldots, m_s)$. The dependencies are visualized in Figure 1. Mathematically, we adopt the linear structural equation model (SEM) with respect to Figure 1

$$\boldsymbol{M} = \boldsymbol{X}\mathbf{U} + \boldsymbol{\varepsilon}^M, \tag{1}$$

$$\boldsymbol{Y} = \boldsymbol{M}\mathbf{V}^\top + \boldsymbol{m}\mathbf{W}^\top + \boldsymbol{\varepsilon}^Y, \tag{2}$$

where $\boldsymbol{\varepsilon}^M \in \mathbb{R}^r$, $\boldsymbol{\varepsilon}^Y \in \mathbb{R}^q$ denote zero-mean errors in the mediators $\boldsymbol{M}$ and traits $\boldsymbol{Y}$ respectively, and $\mathbf{U} \in \mathbb{R}^{p\times r}$, $\mathbf{V} \in \mathbb{R}^{q\times r}$, $\mathbf{W} \in \mathbb{R}^{q\times s}$ are coefficient matrices. We assume the errors $\boldsymbol{\varepsilon}^M$, $\boldsymbol{\varepsilon}^Y$ are independent of the SNPs and the environmental mediators are centered and do not interact with SNPs. Concerning that genetic pathways usually only include a small proportion of SNPs and traits, the coefficient matrices $\mathbf{U}$, $\mathbf{V}$ are often sparse.

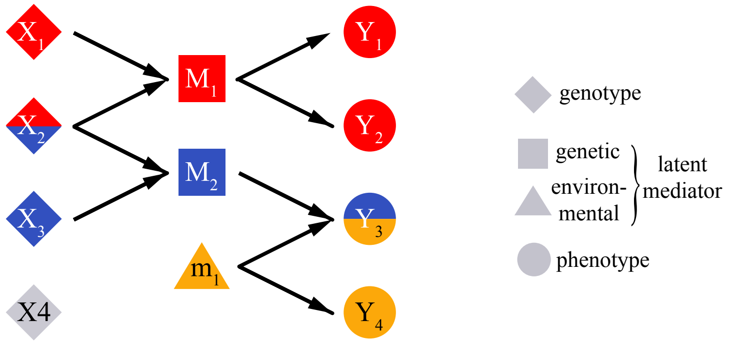

Figure 1: *Dependency of SNPs, traits, latent genetic mediators, and latent environmental mediators. The example contains 4 SNPs and 4 traits. There are 2 latent genetic mediators: genetic mediator $M_1$ is influenced by SNPs $X_1$, $X_2$ and affects traits $Y_1$, $Y_2$; genetic mediator $M_2$ is influenced by SNPs $X_2$, $X_3$ and affects trait $Y_3$. There is one latent environmental mediator $m_1$ which is independent of all SNPs and affects traits $Y_3$, $Y_4$.*

Our goal is to discover genetic pathways: SNPs $\to$ genetic latent mediator $\to$ traits. Since the genetic mediators are unobserved, we look for clusters of SNPs and traits potentially related to the same underlying mediator. For example, there are two genetic pathways in Figure 1: $X_1, X_2 \to M_1 \to Y_1, Y_2$ in red and $X_2, X_3 \to M_2 \to Y_3$ in blue. We aim to uncover the corresponding gene-trait clusters $\{X_1, X_2, Y_1, Y_2\}$ and $\{X_2, X_3, Y_3\}$. Under model (1), (2), let $\mathbf{U}_{\cdot k}$, $\mathbf{V}_{\cdot k}$ be the $k$-th column of $\mathbf{U}$, $\mathbf{V}$, then the $k$-th genetic pathway refers to the SNPs' effects on the $k$-th mediator $M_k = \boldsymbol{X}\mathbf{U}_{\cdot k} + \varepsilon_k^M$ and the mediator's effect on the traits $M_k\mathbf{V}_{\cdot k}^\top$. The $k$-th gene-trait cluster consists of SNPs and traits with non-zero loadings in $\mathbf{U}_{\cdot k}$, $\mathbf{V}_{\cdot k}$.

Our analysis builds upon genome-wide association studies (GWAS) summary statistics due to their wider availability compared to individual-level data $\{(\boldsymbol{X}, \boldsymbol{Y})\}$. In particular, we focus on SNP-trait effect (marginal association) estimates $\hat{\beta}_{ij}$ of SNP-trait pairs $(X_i, Y_j)$ obtained from simple linear regressions. Assume the SNP-trait effects are derived from a sample of size $n$. We denote the matrix of genes of $n$ individuals by $\mathbf{X} \in \mathbb{R}^{n\times p}$, where the $i$-th row $\mathbf{X}_{i\cdot}$ represents the genes of the $i$-th individual, and the $j$-th column $\mathbf{X}_{\cdot j}$ represents the $j$-th SNP. Similarly we define $\mathbf{Y} \in \mathbb{R}^{n\times q}$, $\mathbf{M} \in \mathbb{R}^{n\times r}$, $\mathbf{m} \in \mathbb{R}^{n\times s}$, $\mathbf{E}^M \in \mathbb{R}^{n\times r}$, and $\mathbf{E}^Y \in \mathbb{R}^{n\times p}$. Assuming the columns of $\mathbf{X}$ are centered, the marginal association estimates take the form

$$\hat{\beta}_{ij} = \frac{\mathbf{X}_{\cdot i}^\top \mathbf{Y}_{\cdot j}}{\mathbf{X}_{\cdot i}^\top \mathbf{X}_{\cdot i}}, \quad 1 \le i \le p,\ 1 \le j \le q. \tag{3}$$

We denote the matrix of marginal association estimates by $\hat{\boldsymbol{\beta}} = (\hat{\beta}_{ij}) \in \mathbb{R}^{p\times q}$.

We draw connections between the marginal association estimates $\hat{\boldsymbol{\beta}}$ and coefficient matrices $\mathbf{U}$, $\mathbf{V}$. First, we plug Eq. (1) into Eq. (2) and rewrite the model in the matrix form

$$\mathbf{Y} = \mathbf{X}\mathbf{U}\mathbf{V}^\top + \mathbf{m}\mathbf{W}^\top + \mathbf{E}^M\mathbf{V}^\top + \mathbf{E}^Y =: \mathbf{X}\mathbf{U}\mathbf{V}^\top + \mathbf{E}', \tag{4}$$

where we introduce the composite error $\mathbf{E}'$ consisting of environmental mediators and two random errors. Eq. (4) decomposes the responses $\mathbf{Y}$ into a genetic component $\mathbf{X}\mathbf{U}\mathbf{V}^\top$ and a non-genetic component $\mathbf{E}'$. The genetic component is linear in $\mathbf{X}$ and we regard the coefficient matrix as the true SNP-trait effect

$$\underbrace{\boldsymbol{\beta}}_{p\times q} := \underbrace{\mathbf{U}}_{p\times r}\underbrace{\mathbf{V}^\top}_{r\times q}. \tag{5}$$

The element $\beta_{ij}$ quantifies the $i$-th SNP's effect on the $j$-th trait. Since both $\mathbf{U}$ and $\mathbf{V}$ are sparse and of rank $r \ll p, q$, the resulting matrix product $\boldsymbol{\beta}$ is also sparse and low-rank. Further assume the environmental mediators are centered, then $\mathbf{E}'$ is zero-mean conditional on $\mathbf{X}$ by the independence of SNPs and $\boldsymbol{m}$, $\boldsymbol{\varepsilon}^M$, $\boldsymbol{\varepsilon}^Y$. We use independent SNPs so that $\mathbf{X}_{\cdot i_1}^\top \mathbf{X}_{\cdot i_2}$, $i_1 \neq i_2$, is approximately zero. In this way, Eq. (3) can be simplified as

$$\hat{\beta}_{ij} = \frac{\mathbf{X}_{\cdot i}^\top (\mathbf{X}\mathbf{U}\mathbf{V}_{j\cdot}^\top + \mathbf{E}'_{\cdot j})}{\mathbf{X}_{\cdot i}^\top \mathbf{X}_{\cdot i}} = \frac{\mathbf{X}_{\cdot i}^\top \mathbf{X}_{\cdot i}\mathbf{U}_{i\cdot}\mathbf{V}_{j\cdot}^\top}{\mathbf{X}_{\cdot i}^\top \mathbf{X}_{\cdot i}} + \frac{\mathbf{X}_{\cdot i}^\top \mathbf{E}'_{\cdot j}}{\mathbf{X}_{\cdot i}^\top \mathbf{X}_{\cdot i}} = \mathbf{U}_{i\cdot}\mathbf{V}_{j\cdot}^\top + \mathbf{E}_{ij} = \beta_{ij} + \mathbf{E}_{ij}, \tag{6}$$

where we use $\mathbf{E}_{ij}$ to denote $\mathbf{X}_{\cdot i}^\top \mathbf{E}'_{\cdot j}/\mathbf{X}_{\cdot i}^\top \mathbf{X}_{\cdot i}$. As a result, we expect the marginal association estimate matrix $\hat{\boldsymbol{\beta}}$ to be close to $\boldsymbol{\beta}$, thus approximately low-rank and sparse.

We end the section by introducing definitions and notations used in the following sections. In the GWAS datasets, each marginal association estimate comes with a statistical significance test. We call a SNP a signal SNP if it is statistically significant to at least one trait. We call a SNP a noise SNP if it has no statistically significant association with any traits. We use superscript 0 for quantities associated with noise SNPs. For a matrix $\mathbf{A}$, we denote its column space by $\mathsf{colsp}(\mathbf{A})$ and its Frobenius norm by $\|\mathbf{A}\|_{\mathrm{F}}$. For a random matrix $\mathbf{G} \in \mathbb{R}^{m\times n}$, we write $\mathbf{G} \sim \mathcal{MN}(\boldsymbol{\mu}, \boldsymbol{\Sigma}, \widetilde{\boldsymbol{\Sigma}})$ if $\mathbf{G}$ follows the matrix normal distribution with mean matrix $\boldsymbol{\mu} \in \mathbb{R}^{m\times n}$ and covariance matrices $\boldsymbol{\Sigma} \in \mathbb{R}^{m\times m}$, $\widetilde{\boldsymbol{\Sigma}} \in \mathbb{R}^{n\times n}$. This is equivalent to vectorized $\mathbf{G}$ following the multivariate normal distribution with vectorized $\boldsymbol{\mu}$ as the mean and $\widetilde{\boldsymbol{\Sigma}} \otimes \boldsymbol{\Sigma}$ as the covariance matrix. Here vectorized $\boldsymbol{\mu}$ denotes a vector of length $mn$ and the $((r-1)m+s)$-th element is $\mu_{s,r}$, i.e., stacking the columns of $\boldsymbol{\mu}$ into a single column vector. The notation $\otimes$ denotes the Kronecker product and $\widetilde{\boldsymbol{\Sigma}} \otimes \boldsymbol{\Sigma}$ is a matrix of dimension $mn \times mn$ and satisfies $(\widetilde{\boldsymbol{\Sigma}} \otimes \boldsymbol{\Sigma})_{m(r-1)+v,m(s-1)+w} = \widetilde{\boldsymbol{\Sigma}}_{rs}\boldsymbol{\Sigma}_{vw}$.

### 2 Method

In this section, we discuss how to reconstruct gene-trait clusters from marginal association estimates $\hat{\boldsymbol{\beta}}$. We first introduce the general method, and then discuss a technique to stabilize the results via bootstrap aggregation (bagging).

#### 2.1 Pipeline

Notice that without further assumptions, matrices $\mathbf{U}$, $\mathbf{V}$ in Eq. (5) are not identifiable even if the true $\boldsymbol{\beta}$ is known[1]. However, $\mathsf{colsp}(\mathbf{U})$ and $\mathsf{colsp}(\mathbf{V})$ are uniquely determined by $\boldsymbol{\beta}$. Therefore, we start with estimating the column spaces in Section 2.1.1. We remark that $\mathsf{colsp}(\mathbf{U})$ and $\mathsf{colsp}(\mathbf{V})$ are of dimension $r$, usually much smaller than the numbers of SNPs and traits. Based on the column space estimators $\widehat{\mathsf{colsp}}(\mathbf{U})$ and $\widehat{\mathsf{colsp}}(\mathbf{V})$, in Section 2.1.2, we propose methods to find $\hat{\mathbf{U}}$, $\hat{\mathbf{V}}$ whose columns are sparse and form bases of the estimated column spaces $\widehat{\mathsf{colsp}}(\mathbf{U})$, $\widehat{\mathsf{colsp}}(\mathbf{V})$, respectively. Finally, we construct a gene-trait cluster for each column pair $(\hat{\mathbf{U}}_{\cdot k}, \hat{\mathbf{V}}_{\cdot k})$, $1 \leq k \leq r$, which estimates the $k$-th genetic pathway. The whole procedure is summarized in Algorithm 1.

---

**Algorithm 1:** PathGPS

---

**Input**: Marginal association estimate matrices of signal SNPs $\hat{\boldsymbol{\beta}}$ and noise SNPs $\hat{\boldsymbol{\beta}}^0$, number of latent mediators $r$.

**Initialization**: Gene-trait cluster list $\mathcal{L} = \emptyset$.

1. Let the number of signal and noise SNPs be $p$, $p^0$, respectively. Compute the truncated eigen-decomposition with $r$ components $\tilde{\mathbf{V}}\tilde{\mathbf{D}}\tilde{\mathbf{V}}^\top$ of

$$\hat{\boldsymbol{\beta}}^\top \hat{\boldsymbol{\beta}} - \frac{p}{p^0} \hat{\boldsymbol{\beta}}^{0^\top} \hat{\boldsymbol{\beta}}^0,$$

where $\tilde{\mathbf{V}} \in \mathbb{R}^{q \times r}$ is orthonormal, $\tilde{\mathbf{D}} \in \mathbb{R}^{r \times r}$ is diagonal. This is a denoising step to remove the environmental confounders. Let $\tilde{\mathbf{U}} = \hat{\boldsymbol{\beta}}\tilde{\mathbf{V}}$.

2. Apply Varimax or Promax to $\tilde{\mathbf{V}}$ and find the transformation matrix $\mathbf{R} \in \mathbb{R}^{r \times r}$ inducing sparse loadings. Let $\hat{\mathbf{V}} = \tilde{\mathbf{V}}\mathbf{R}$ , $\hat{\mathbf{U}} = \tilde{\mathbf{U}}(\mathbf{R}^{-1})^\top$.

3. **for** $k$ *from* 1 *to* $r$ **do**

   Define the $k$-th gene-trait cluster as

$$C_k := \left\{X_i : \hat{\mathbf{U}}_{ik} \neq 0\right\} \cup \left\{Y_j : \hat{\mathbf{V}}_{jk} \neq 0\right\}.$$

   Add the $k$-th cluster into the cluster list $\mathcal{L} \leftarrow \mathcal{L} \cup \{C_k\}$.

**end**

**Output**: Gene-trait cluster list $\mathcal{L}$.

---

##### 2.1.1 Column space estimation

Provided with the true number of latent mediators $r$, arguably the most straightforward approach of column space estimation is

[1] In fact, for any invertible matrix $\mathbf{R} \in \mathbb{R}^{r \times r}$, define $\mathbf{V}' = \mathbf{V}\mathbf{R}$, $\mathbf{U}' = \mathbf{U}\left(\mathbf{R}^{-1}\right)^\top$, then $\mathbf{U}'\mathbf{V}'^\top = \mathbf{U}\mathbf{V}^\top = \boldsymbol{\beta}$.

1. Compute the singular value decomposition (SVD) of $\hat{\boldsymbol{\beta}}$ and keep the top $r$ left and right singular vectors $\hat{\mathbf{U}}_{\text{SVD}}$, $\hat{\mathbf{V}}_{\text{SVD}}$;
2. Let $\widehat{\mathsf{colsp}}(\mathbf{U}) = \mathsf{colsp}(\hat{\mathbf{U}}_{\text{SVD}})$, $\widehat{\mathsf{colsp}}(\mathbf{V}) = \mathsf{colsp}(\hat{\mathbf{V}}_{\text{SVD}})$.

However, as shown below, the marginal associations $\hat{\boldsymbol{\beta}}$ could be a mixture of genetic and environmental influences. The simple SVD is unable to tell apart the environmental influences from the genetic components.

**Proposition 1.** *Under model (4) with $n$ independent individuals and $p$ SNPs, assume*

1. *Given $\boldsymbol{X}$, random vectors/variables $\boldsymbol{m}$, $\boldsymbol{\varepsilon}^M$, $\boldsymbol{\varepsilon}^Y$ are with zero means, covariance matrices $\boldsymbol{\Sigma}_m$, $\boldsymbol{\Sigma}_{\boldsymbol{\varepsilon}^M}$, $\boldsymbol{\Sigma}_{\boldsymbol{\varepsilon}^Y}$, and uncorrelated;*
2. $\mathbf{X}^\top \mathbf{X} = n\mathbf{I}_p$.

*Then the marginal association estimates satisfy*

$$\mathbb{E}\left[\hat{\boldsymbol{\beta}}^\top \hat{\boldsymbol{\beta}}\right] = \mathbf{V}\mathbf{U}^\top \mathbf{U}\mathbf{V}^\top + \frac{p}{n}\left(\mathbf{W}\boldsymbol{\Sigma}_m\mathbf{W}^\top + \mathbf{V}\boldsymbol{\Sigma}_{\boldsymbol{\varepsilon}^M}\mathbf{V}^\top + \boldsymbol{\Sigma}_{\boldsymbol{\varepsilon}^Y}\right), \tag{7}$$

*where the expectation is conditional on $\mathbf{X}$.*

Proposition 1 suggests that the column space of $\hat{\boldsymbol{\beta}}^\top \hat{\boldsymbol{\beta}} \approx \mathbb{E}\left[\hat{\boldsymbol{\beta}}^\top \hat{\boldsymbol{\beta}}\right]$ is contaminated by the environmental covariance $\mathbf{W}\boldsymbol{\Sigma}_m\mathbf{W}^\top$ and the response error covariance $\boldsymbol{\Sigma}_{\boldsymbol{\varepsilon}^Y}$, especially when the ratio $p/n$ is not ignorable. Thus, the simple SVD may mistakenly regard the non-genetic influences as part of $\mathsf{colsp}(\mathbf{V})$. Figure 2 panel (a) displays a simulation study where the simple SVD approach (in dark blue) fails in the presence of environmental factors.

To separate the genetic and environmental components in Eq. (7), we propose a method using noise SNPs. The idea is to use the marginal association estimates $\hat{\boldsymbol{\beta}}^0$ of noise SNPs to estimate the non-genetic structure and remove it from the marginal association estimates $\hat{\boldsymbol{\beta}}$ of signal SNPs.

The following proposition is the counterpart of Proposition 1 for noise SNPs.

**Proposition 2.** *Under model (4) with $n$ independent individuals, assumptions in Proposition 1, and further assume*

1. ${\mathbf{X}^0}^\top \mathbf{X}^0 = n\mathbf{I}_{p^0}$, ${\mathbf{X}^0}^\top \mathbf{X} = \mathbf{0}$.

*Then the marginal association estimates of noise SNPs satisfy*

$$\mathbb{E}\left[{\hat{\boldsymbol{\beta}}^0}^\top \hat{\boldsymbol{\beta}}^0\right] = \frac{p^0}{n}\left(\mathbf{W}\boldsymbol{\Sigma}_m\mathbf{W}^\top + \mathbf{V}\boldsymbol{\Sigma}_{\boldsymbol{\varepsilon}^M}\mathbf{V}^\top + \boldsymbol{\Sigma}_{\boldsymbol{\varepsilon}^Y}\right), \tag{8}$$

*where the expectation is conditional on $\mathbf{X}$ and $\mathbf{X}^0$.*

The proposed method relies on the following corollary of Proposition 1 and Proposition 2.

**Corollary 1.** *Under model (4) with $n$ independent individuals, assumptions in Proposition 1 and Proposition 2, the marginal association estimates satisfy*

$$\mathbb{E}\left[\hat{\boldsymbol{\beta}}^\top \hat{\boldsymbol{\beta}} - \frac{p}{p^0}{\hat{\boldsymbol{\beta}}^0}^\top \hat{\boldsymbol{\beta}}^0\right] = \mathbf{V}\mathbf{U}^\top \mathbf{U}\mathbf{V}^\top, \tag{9}$$

*where the expectation is conditional on $\mathbf{X}$ and $\mathbf{X}^0$.*

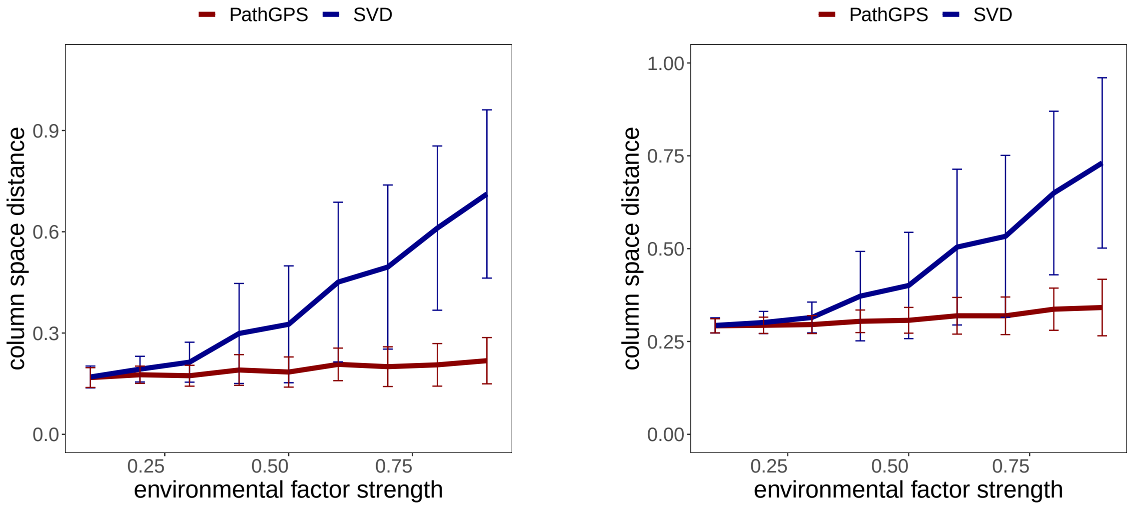

(a) colsp($\mathbf{V}$)  (b) colsp($\mathbf{U}$)

Figure 2: *Column space estimation. We compare the simple SVD approach and the proposed subtraction estimator. The data generation mechanism and the column space distance are discussed in Section 3. We plot the average column space distances for* $\mathbf{V}$ *(panel (a)) and* $\mathbf{U}$ *(panel (b)), plus and minus one standard deviation. All results are aggregated over* 100 *trials.*

It is desirable to use data from multiple cohorts, but those cohorts may come from different populations and measure different traits. We show that Corollary 1 is valid with multiple populations.

**Corollary 2.** *Assume that $K$ independent populations follow model (1), (2) and satisfy the assumptions in Proposition 1 and 2. Suppose that each marginal association is computed based on a subset of the $K$ populations, then*

$$\mathbb{E}\left[\hat{\boldsymbol{\beta}}^\top \hat{\boldsymbol{\beta}} - \frac{p}{p_0}\hat{\boldsymbol{\beta}}^{0\top}\hat{\boldsymbol{\beta}}^0\right] = \mathbf{V}\mathbf{U}^\top\mathbf{U}\mathbf{V}^\top,$$

*where the expectation is conditional on* $\mathbf{X}$ *and* $\mathbf{X}^0$.

Motivated by Corollary 1 and 2, we introduce the subtraction estimator of colsp($\mathbf{V}$):

1. Compute the truncated eigen-decomposition with $r$ components

$$\hat{\boldsymbol{\beta}}^\top \hat{\boldsymbol{\beta}} - \frac{p}{p^0}\hat{\boldsymbol{\beta}}^{0^\top}\hat{\boldsymbol{\beta}}^0 \approx \tilde{\mathbf{V}}\tilde{\mathbf{D}}\tilde{\mathbf{V}}^\top. \tag{10}$$

2. Let $\widehat{\mathsf{colsp}}(\mathbf{V}) = \mathsf{colsp}(\tilde{\mathbf{V}})$.

Since the environmental influences are removed by subtracting a scalar multiple of $\hat{\boldsymbol{\beta}}^{0^\top}\hat{\boldsymbol{\beta}}^0$, the estimated column spaces should consist solely of the genetic components. In Figure 2 panel (a), we observe the proposed method (in dark red) is robust to the environmental factors in the simulation study.

As for $\mathsf{colsp}(\mathbf{U})$, by Eq. (15) in Section 4, the first term $\mathbf{U}\mathbf{V}^\top$ in $\hat{\boldsymbol{\beta}}$ represents the genetic signal and contains information of $\mathsf{colsp}(\mathbf{U})$; the second term $(\mathbf{X}^\top \mathbf{m}\mathbf{W}^\top + \mathbf{X}^\top \mathbf{E}^M \mathbf{V}^\top + \mathbf{X}^\top \mathbf{E}^Y)/n = \mathbf{E}$ by Eq. (6) represents the non-genetic noise and is irrelevant to $\mathsf{colsp}(\mathbf{U})$. Instead of directly analyzing $\hat{\boldsymbol{\beta}}$, we suggest working with $\hat{\boldsymbol{\beta}}\mathbf{V}_\perp$, where $\mathbf{V}_\perp$ is an orthonormal basis of $\mathsf{colsp}(\mathbf{V})$. According to Eq. (6),

$$\hat{\boldsymbol{\beta}}\mathbf{V}_\perp = (\mathbf{U}\mathbf{V}^\top)\mathbf{V}_\perp + \mathbf{E}\mathbf{V}_\perp. \tag{11}$$

After the transformation from $\hat{\boldsymbol{\beta}}$ to $\hat{\boldsymbol{\beta}}\mathbf{V}_\perp$, the Frobenius norm of the genetic signal $\mathbf{U}\mathbf{V}^\top$ is unchanged,

$$\|\mathbf{U}\mathbf{V}^\top \mathbf{V}_\perp\|_{\mathrm{F}}^2 = \mathrm{tr}\left(\mathbf{U}\mathbf{V}^\top \mathbf{V}_\perp \mathbf{V}_\perp^\top \mathbf{V}\mathbf{U}^\top\right) = \mathrm{tr}\left(\mathbf{U}\mathbf{V}^\top \mathbf{V}\mathbf{U}^\top\right) = \|\mathbf{U}\mathbf{V}^\top\|_{\mathrm{F}}^2.$$

Here we use the fact that $\mathbf{V}_\perp \mathbf{V}_\perp^\top$ is the projection operator onto $\mathsf{colsp}(\mathbf{V})$, and $\mathbf{V}_\perp \mathbf{V}_\perp^\top \mathbf{V} = \mathbf{V}$ because $\mathbf{V} \in \mathsf{colsp}(\mathbf{V})$. In contrast, the non-genetic noise is likely to shrink regarding the Frobenius norm again because $\mathbf{V}_\perp \mathbf{V}_\perp^\top$ is a projection operator,

$$\|\mathbf{E}\mathbf{V}_\perp\|_{\mathrm{F}}^2 = \mathrm{tr}\left(\mathbf{E}\mathbf{V}_\perp \mathbf{V}_\perp^\top \mathbf{E}^\top\right) \leq \mathrm{tr}\left(\mathbf{E}\mathbf{E}^\top\right) = \|\mathbf{E}\|_{\mathrm{F}}^2.$$

The equality is obtained only if all rows of $\mathbf{E}$ fall into $\mathsf{colsp}(\mathbf{V})$. As a result, the relative magnitude of the signal over the noise in $\hat{\boldsymbol{\beta}}\mathbf{V}_\perp$ relating to $\mathsf{colsp}(\mathbf{U})$ is increased from that in $\hat{\boldsymbol{\beta}}$. We remark that the property is valid if there are multiple cohorts. When $\mathbf{V}_\perp$ is not available, we replace it by the estimator $\tilde{\mathbf{V}}$ in Eq. (10). Our final estimator of $\mathsf{colsp}(\mathbf{U})$ is

$$\widehat{\mathsf{colsp}}(\mathbf{U}) = \mathsf{colsp}(\hat{\boldsymbol{\beta}}\tilde{\mathbf{V}}). \tag{12}$$

Figure 2 panel (b) compares the simple SVD estimator of $\mathsf{colsp}(\mathbf{U})$ and the proposed estimator (12) in the simulation study. The performance of the naive estimator deteriorates as the environmental influence increases, while the proposed estimator is stable. In addition, we notice an interesting phenomenon of independent interest that the estimators of $\mathsf{colsp}(\mathbf{V})$ are relatively more accurate compared to their counterparts of $\mathsf{colsp}(\mathbf{U})$. We briefly explain the reasons. For the naive approach, by Eq. (15) in Section 4, the error term, in particular $\mathbf{X}^\top \mathbf{E}^M \mathbf{V}^\top/n$ attributed to the genetic mediator noise, contains information about $\mathsf{colsp}(\mathbf{V})$ but not $\mathsf{colsp}(\mathbf{U})$. For the proposed approach, the estimator of $\mathsf{colsp}(\mathbf{U})$ relies on the estimator $\tilde{\mathbf{V}}$ and implicitly $\widehat{\mathsf{colsp}}(\mathbf{V})$, therefore $\widehat{\mathsf{colsp}}(\mathbf{U})$ can not be more precise than $\widehat{\mathsf{colsp}}(\mathbf{V})$.

###### 2.1.2 Matrix rotation and clustering

In this section, we first find sparse matrices in the estimated column spaces. We then construct gene-trait clusters from the non-zero loadings.

Let $\tilde{\mathbf{U}}$, $\tilde{\mathbf{V}}$ be two candidate matrices from th e target column spaces. We aim to find the transformation matrix $\mathbf{R} \in \mathbb{R}^{r\times r}$ such that $\hat{\mathbf{U}} := \tilde{\mathbf{U}}(\mathbf{R}^{-1})^\top$, $\hat{\mathbf{V}} := \tilde{\mathbf{V}}\mathbf{R}$ have sparse columns. We remark that we restrict the matrix product $\tilde{\mathbf{U}}\tilde{\mathbf{V}}^\top$ to be invariant after the transformation for the sake of (5).

There are a number of readily available methods from factor analysis to achieve sparsity. In particular, we discuss two commonly used approaches.

- *Varmiax* [Kai58]. We start from an orthonormal matrix $\tilde{\mathbf{V}}$ and solve

$$\max_{\mathbf{R}^\top\mathbf{R}=\mathbf{R}\mathbf{R}^\top=\boldsymbol{I}_r} \sum_{j=1}^{r} \left( \frac{1}{q}\sum_{i=1}^{q} (\tilde{\mathbf{V}}^\top\mathbf{R})_{ij}^4 - \left( \frac{1}{q}\sum_{i=1}^{q} (\tilde{\mathbf{V}}^\top\mathbf{R})_{ij}^2 \right)^2 \right) \tag{13}$$

  for a rotation matrix $\mathbf{R}$. The criterion (13) denotes the variances of squared loadings of $\tilde{\mathbf{V}}^\top\mathbf{R}$'s columns. The resulting $\hat{\mathbf{V}}$ tend to have many small loadings and we can set those values to zero. Finally, we let $\hat{\mathbf{U}} = \tilde{\mathbf{U}}(\mathbf{R}^{-1})^\top = \tilde{\mathbf{U}}\mathbf{R}$ and set small values in $\hat{\mathbf{U}}$ to zero.

- *Promax* [HW64]. The approach relaxes the orthonormal restriction of $\mathbf{R}$ in Varimax. Promax first applies the above Varimax to $\tilde{\mathbf{V}}$, and then rotates the orthogonal results to a least squares fit. Consequently, the loadings in $\hat{\mathbf{V}}$ are pushed further apart. Again, we can let $\hat{\mathbf{U}} = \tilde{\mathbf{U}}(\mathbf{R}^{-1})^\top$. Similar to *Varimax*, we set small values in $\hat{\mathbf{U}}$, $\hat{\mathbf{V}}$ to zero.

At last, with sparse column estimators $\hat{\mathbf{U}}$, $\hat{\mathbf{V}}$, we loop over $r$ column pairs $(\hat{\mathbf{U}}_{\cdot k}, \hat{\mathbf{V}}_{\cdot k})$ and collect the traits and genes with non-zero loadings.

#### 2.2 Bootstrap aggregation

In this section, we first discuss several issues that may undermine the reliability of Algorithm (1). We then posit a bootstrap aggregation (bagging) approach to stabilize the pipeline and make the results more replicable.

##### 2.2.1 Causes of instability

We discuss three sources of instability.

- *Trait collection.* In modern biobanks, the set of measured traits is evergrowing. Many traits are repeatedly measured in slightly different ways (e.g., disease diagnosis and taking certain drugs that treat the disease). It is desirable to obtain consistent results if we have a slightly different set of traits.

- *Signal SNPs extraction.* In the data preprocessing procedures, we use an external SNP dataset to select uncorrelated index SNPs. Among the chosen index SNPs, we further determine signal and noise SNPs based on the p-values from marginal association tests. We expect to arrive at similar results if we perturb the index SNP sets, especially the signal SNPs, by a small amount.

- *Hyper-parameter selection.* The pipeline relies on the number of latent mediators $r$, the matrix transformation method, and the number of non-zero loadings preserved after the matrix transformation. The cluster list $\mathcal{L}$ should be robust to hyper-parameter selection.

##### 2.2.2 Co-appearance graph and SNP bootstrapping

To stabilize Algorithm 2, we propose to perturb the entire procedure many times and then aggregate the results. We first introduce our aggregation method via a co-appearance graph, and then delve into the perturbation: SNP bootstrapping.

**Definition 1** (Co-appearance frequency)**.** *Let $\mathcal{A}$ be a set of interested elements, $C_k$ be clusters defined on $\mathcal{A}$, and $\mathcal{L}_b = \{C_k\}$ be a list of such clusters. For each pair of elements $a_i$, $a_j \in \mathcal{A}$, we define the co-appearance frequency as*

$$w_{ij} = \frac{1}{B} \sum_{b=1}^{B} \frac{1}{|\mathcal{L}_b|} \sum_{C_k \in \mathcal{L}_b} \mathbb{1}_{\{i \in C_k\}} \mathbb{1}_{\{j \in C_k\}}, \tag{14}$$

*where $|\mathcal{L}_b|$ denotes the number of clusters in the list $\mathcal{L}_b$.*

The co-appearance frequency (14) describes how frequently two elements appear in the same cluster. If two elements always show up in the same cluster, the pair will have a high co-appearance frequency, and we are more confident that the pair are related. In our application, we are interested in the element set including all candidate signal SNPs and traits. We define the co-appearance frequency for gene-gene, gene-trait, and trait-trait pairs as Eq. (14).

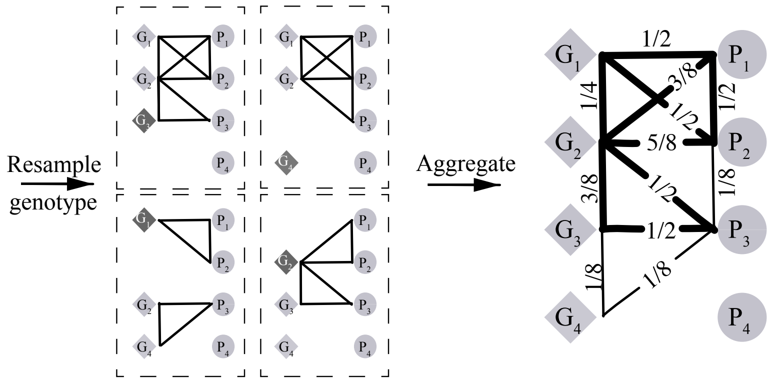

Figure 3: *SNP bootstrapping and co-appearance graph.*

With all the co-appearance frequencies, we can construct a weighted undirected graph of genes and traits. We let each node represent a gene or a trait, and weight each edge $(i, j)$ by $w_{ij}$ in Eq. (14). Figure 3 displays an example. The weighted graph is convenient for downstream analysis. In particular, we focus on two threads of clustering methods.

- *Dimension reduction and clustering.* We use t-SNE [HR02] or UMAP [MHM18] to find low-dimensional embeddings. The embeddings can be used to visualize genes and traits that are closely connected in the co-appearance graph. The representations can also be fed to various clustering methods based on feature vectors, such as k-means.

- *Graph clustering.* There are a multitude of graph clustering methods available, such as spectral clustering and label propagation.

Now we discuss the SNP bootstrapping for generating cluster lists. In each trial, we resample the same number of signal SNPs with replacement and obtain bootstrapped signal marginal association estimates $\hat{\boldsymbol{\beta}}^b$. We then apply Algorithm 1 to $\hat{\boldsymbol{\beta}}^b$ and $\hat{\boldsymbol{\beta}}^0$ and arrive at a cluster list $\mathcal{L}_b$. We repeat the trial $B$ times and get a collection of cluster lists $\{\mathcal{L}_b\}$. Figure 3 describes the bootstrap process. Algorithm 2 summarizes the complete bagging procedure. We remark that the column estimator $\hat{\mathbf{U}}$ may have duplicate rows due to sampling with replacement, but we merge different copies of the same signal SNP in computing the co-appearance frequency (14).

---

**Algorithm 2:** Bagged PathGPS

---

**Input**: Marginal association estimate matrix of signal SNPs $\hat{\boldsymbol{\beta}}$, marginal association estimate matrix of noise SNPs $\hat{\boldsymbol{\beta}}^0$, number of latent mediators $r$, number of bootstrap trials $B$.

**Initialization**: A set of gene-trait cluster list $\mathcal{S} = \emptyset$.

1.**for** $b$ *from* 1 *to* $B$ **do**

a. Resample the same number of signal genes with replacement and obtain $\hat{\boldsymbol{\beta}}^b$.

b. Apply Algorithm 1 with inputs $\hat{\boldsymbol{\beta}}^b$, $\hat{\boldsymbol{\beta}}^0$, $r$, and get a cluster list $\mathcal{L}_b$. Update the set of lists $\mathcal{S} \leftarrow \mathcal{S} \cup \{\mathcal{L}_b\}$.

**end**

2. Compute the co-appearance frequency (14) using $\mathcal{S}$ for all signal SNPs and traits.

3. Use t-SNE/UMAP to find low-dimensional embeddings and cluster the embedded coordinates. Or run graph clustering algorithms on the co-appearance graph to get gene-trait clusters. Denote the final list of clusters by $\mathcal{L}$.

**Output**: Gene-trait cluster list $\mathcal{L}$.

---

We discuss how bagging (Algorithm 2) helps with the three sources of instability in Section 2.2.1.

- *Trait collection.* In Section 6, we subsample traits without replacement and evaluate the stability of clusters. Regardless of the clustering methods, the bootstrap aggregation produces more stable clusters compared to the one-shot counterpart.

- *Signal SNPs extraction.* By the design of SNP bootstrapping, the aggregated method is robust to perturbations of signal SNPs. Therefore, with a different but largely overlapped set of signal SNPs, the aggregated method should yield similar clusters.

- *Hyper-parameter selection.* In Figure 4, we empirically compare three versions of our method: one-shot pipeline, clustering based on the co-appearance graph from one-shot pipeline, and clustering based on the co-appearance graph with bootstrapping. We test the robustness of the methods when the hyper-parameters—the latent genetic mediator numbers $r$ and the number of genes and traits per cluster $n_\emptyset$—are misspecified. Across different simulation settings, the bootstrap aggregated approach are in general the most robust. We provide intuitions of why the bagging improves the stability.

  - $n'_\emptyset > n_\emptyset$ *or* $r' > r$. When we include more genes or traits in a cluster, i.e., $n'_\emptyset > n_\emptyset$, SNP-trait pairs which do not share any genetic pathways may be selected into a cluster. However, pairs with no biological connection are chosen randomly, while pairs sharing biological pathways are selected almost surely. Therefore, the co-appearance frequencies $w_{ij}$ of SNP-trait pairs which do not share any genetic pathways should be smaller than those of pairs that share. Similarly for $r' > r$.

– $n'_{\emptyset} < n_{\emptyset}$ *or* $r' < r$. When we include less genes or traits in a cluster, i.e., $n'_{\emptyset} < n_{\emptyset}$, there will inevitably be false negatives. However, in the bagging approach, we perturb the signal SNPs each time and the true non-zero loadings are likely to be selected in a certain proportion of the bootstrap estimates. Similarly for $r' < r$.

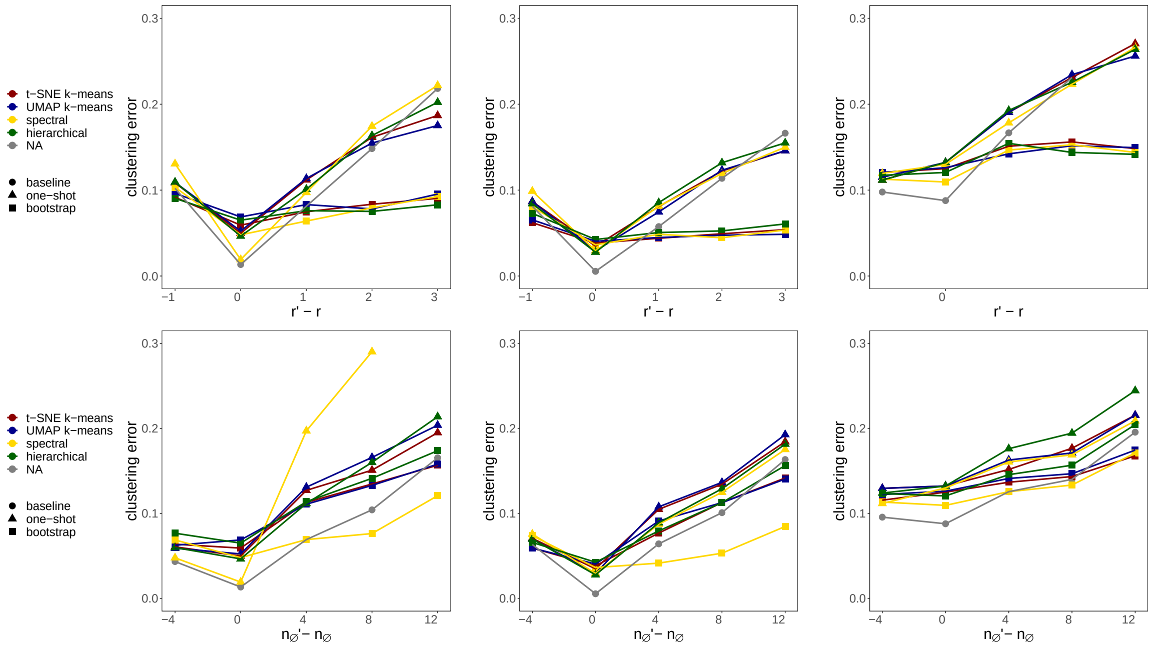

*(a) Default* *(b) Sparse* *(c) Overlap*

Figure 4: *Robustness to hyper-parameter misspecification. We compare methods with and without bagging. Data generation mechanisms, method details, and the clustering error are discussed in Section 3. We input a sequence of $r'$ (first row) and $n'_{\emptyset}$ (second row) around the true values. We plot the average clustering errors plus and minus one standard deviation. All results are aggregated over* 100 *trials.*

### 3 Simulation

#### 3.1 Column space estimation

We describe the simulation settings of Figure 2 comparing the simple SVD approach and the proposed subtraction estimator.

We generate data following the hierarchical model (1), (2). We consider $n = 2000$ individuals, $p = 100$ signal SNPs, $p^0 = 400$ noise SNPs, $q = 100$ traits, $r = 4$ latent genetic mediators, and $s = 2$ latent environmental mediators. We generate signal/noise SNPs and latent environmental mediators independently from standard Gaussian distribution. As for coefficient matrices, we generate $\mathbf{U}$, $\mathbf{V}$'s elements uniformly from $[-1, 1]$, and randomly set 80% of the elements to zero. We generate elements of the environmental mediators' coefficient matrix $\mathbf{W}$ uniformly from $[-c, c]$, random errors $\boldsymbol{\varepsilon}^M$, $\boldsymbol{\varepsilon}^Y$ from Gaussian distribution, and adjust $c$, magnitudes of

$\varepsilon^M$, $\varepsilon^Y$ so that the variance of the composite error $\varepsilon' = \boldsymbol{m}\mathbf{W} + \varepsilon^M \mathbf{V}^\top + \varepsilon^Y$ stays the same but the proportion of the environmental mediators' variance $\mathsf{Var}(\boldsymbol{m}\mathbf{W})/\mathsf{Var}(\varepsilon')$ (environmental factor strength) varies from 10% to 90%.

We measure the performance of column space estimators by the following linear space distance. For two linear spaces spanned by the columns of $\mathbf{U}_1$, $\mathbf{U}_2$, respectively, we define the distance

$$\mathrm{dist}(\mathsf{colsp}(\mathbf{U}_1), \mathsf{colsp}(\mathbf{U}_2)) := \max_{\boldsymbol{\xi} \in \mathbb{R}^p, \|\boldsymbol{\xi}\|_2 = 1} \|\boldsymbol{P}_{\mathbf{U}_1}\boldsymbol{\xi} - \boldsymbol{P}_{\mathbf{U}_2}\boldsymbol{\xi}\|_2,$$

where $\boldsymbol{P}_{\mathbf{U}_1}$, $\boldsymbol{P}_{\mathbf{U}_2}$ are the projection operators onto the column spaces of $\mathbf{U}_1$, $\mathbf{U}_2$.

##### 3.2 Robustness to hyper-parameter misspecification

We describe the simulation settings of Figure 4 comparing methods with and without bagging.

The default setting follows the data generation mechanism in Figure 2 with $n = 500$, $p = 50$, $p^0 = 150$, $q = 30$, $r = 3$, $s = 2$, and a total of $n_\emptyset = 16$ genes and traits per cluster. We consider two variations: a sparser setting with less genes and traits per cluster and an overlap setting where a gene/trait may belong to multiple clusters. Details are summarized in Table 1.

| Setting | total number of genes and traits/cluster ($n_\emptyset$) | multiple membership |
|---|---|---|
| Default | 16 | × |
| Sparse | 12 | × |
| Overlap | 16 | ✓ |

Table 1: *Summary of simulation settings of Figure 4.*

In each setting, we input a sequence of $r'$ and $n'_\emptyset$ around the true values to three versions of our method: the one-shot pipeline (baseline) follows Algorithm 1 (no further clustering is required, clustering method denoted by "NA" in the legend), the clustering without bootstrapping (one-shot) forms a co-appearance graph of the list from the one-shot pipeline and applies various clustering methods, and the co-appearance clustering with bootstrapping (bootstrap) implements Algorithm 1 with 200 bootstrap resamples. As for clustering methods, we consider two methods with low-dimensional embeddings: t-SNE and UMAP with k-means, and two graph clustering methods: spectral clustering and hierarchical clustering.

We evaluate the performance of the clustering by the minimal clustering error across label permutation below. In particular, let $\mathcal{L}$ be the true list of $r$ clusters defined on set $\mathcal{A}$, and $\hat{\mathcal{L}}$ be an estimate with the same number of clusters , then we define the clustering error

$$\min_{\pi} \frac{1}{r} \sum_{k=1}^{r} \frac{1}{|\mathcal{A}|} \sum_{a \in \mathcal{A}} \left( \mathbb{1}_{\{a \in C_{\pi(k)}, a \notin \hat{C}_k\}} + \mathbb{1}_{\{a \notin C_{\pi(k)}, a \in \hat{C}_k\}} \right),$$

where $\pi : \{1, 2, \ldots, r\} \to \{1, 2, \ldots, r\}$ denotes permutation over cluster labels.

#### 4 Proof

*Proof of Proposition 1.* Notice that if $\mathbf{X}^\top \mathbf{X}$ is diagonal, the marginal association estimates and the coefficients from the full linear regression of all SNPs are equivalent. For simplicity, under

Assumption 2, we stick to the latter

$$\begin{aligned}\hat{\boldsymbol{\beta}} &= (\mathbf{X}^\top \mathbf{X})^{-1}\mathbf{X}^\top \mathbf{Y} = \frac{\boldsymbol{X}^\top \boldsymbol{Y}}{n} \\ &= \mathbf{U}\mathbf{V}^\top + \frac{\mathbf{X}^\top \mathbf{m}\mathbf{W}^\top}{n} + \frac{\mathbf{X}^\top \mathbf{E}^M \mathbf{V}^\top}{n} + \frac{\mathbf{X}^\top \mathbf{E}^Y}{n}.\end{aligned} \tag{15}$$

Since Eq. (7) only depends on the first and second order moments of $\boldsymbol{m}$, $\boldsymbol{\varepsilon}^M$, $\boldsymbol{\varepsilon}^Y$, we can prove the statement assuming

$$\mathbf{m} \sim \mathcal{MN}(\mathbf{0}, \mathbf{I}_n, \boldsymbol{\Sigma}_m), \quad \mathbf{E}^M \sim \mathcal{MN}(\mathbf{0}, \mathbf{I}_n, \boldsymbol{\Sigma}_{\boldsymbol{\varepsilon}^M}), \quad \mathbf{E}^Y \sim \mathcal{MN}(\mathbf{0}, \mathbf{I}_n, \boldsymbol{\Sigma}_{\boldsymbol{\varepsilon}^Y}), \tag{16}$$

and $\boldsymbol{m}$, $\boldsymbol{\varepsilon}^M$, $\boldsymbol{\varepsilon}^Y$ are independent.

Now we derive (7). Based on (15) and Assumption 1, we compute the expectation of $\hat{\boldsymbol{\beta}}^\top \hat{\boldsymbol{\beta}}$ and omit all the zero-expectation interaction terms involving $\mathbf{m}$, $\mathbf{E}^M$, $\mathbf{E}^Y$,

$$\begin{aligned}\mathbb{E}\left[\hat{\boldsymbol{\beta}}^\top \hat{\boldsymbol{\beta}}\right] =& \mathbf{V}\mathbf{U}^\top \mathbf{U}\mathbf{V}^\top + \mathbb{E}\left[\frac{\mathbf{W}\mathbf{m}^\top \mathbf{X}}{n}\frac{\mathbf{X}^\top \mathbf{m}\mathbf{W}^\top}{n}\right] \\ &+ \mathbb{E}\left[\frac{\mathbf{V}{\mathbf{E}^M}^\top \mathbf{X}}{n}\frac{\mathbf{X}^\top \mathbf{E}^M \mathbf{V}^\top}{n}\right] + \mathbb{E}\left[\frac{{\mathbf{E}^Y}^\top \mathbf{X}}{n}\frac{\mathbf{X}^\top \mathbf{E}^Y}{n}\right].\end{aligned} \tag{17}$$

According to [GN18, Chapter 2], we have the following properties of matrix normal distribution.

**Lemma 1.**

1. *Let* $\mathbf{R}$ *be a matrix of independent standard normals,* $\mathbf{M}$*,* $\mathbf{N}$ *are matrices, and let* $\mathbf{H} = \mathbf{M}\mathbf{R}\mathbf{N}$*, then*

$$\mathbf{H} \sim \mathcal{MN}(\mathbf{0}, \mathbf{M}\mathbf{M}^\top, \mathbf{N}^\top \mathbf{N}).$$

2. *Let* $\mathbf{R} \sim \mathcal{MN}(\mathbf{0}, \mathbf{M}, \mathbf{N})$*, then*

$$\mathbb{E}[\mathbf{R}\mathbf{R}^\top] = \mathrm{tr}(\mathbf{N})\mathbf{M}, \quad \mathbb{E}[\mathbf{R}^\top \mathbf{R}] = \mathrm{tr}(\mathbf{M})\mathbf{N}.$$

Apply Lemma 1 to the random matrices in Eq. (17),

$$\begin{aligned}&\frac{\mathbf{W}\mathbf{m}^\top \mathbf{X}}{\sqrt{n}} \sim \mathcal{MN}(\mathbf{0}, \mathbf{W}\boldsymbol{\Sigma}_m \mathbf{W}^\top, \mathbf{I}_p), \quad \mathbb{E}\left[\frac{\mathbf{W}\mathbf{m}^\top \mathbf{X}}{\sqrt{n}}\left(\frac{\mathbf{W}\mathbf{m}^\top \mathbf{X}}{\sqrt{n}}\right)^\top\right] = \mathrm{tr}(\mathbf{I}_p)\mathbf{W}\boldsymbol{\Sigma}_m \mathbf{W}^\top, \\ &\frac{\mathbf{V}{\mathbf{E}^M}^\top \mathbf{X}}{\sqrt{n}} \sim \mathcal{MN}(\mathbf{0}, \mathbf{V}\boldsymbol{\Sigma}_{\boldsymbol{\varepsilon}^M}\mathbf{V}^\top, \mathbf{I}_p), \quad \mathbb{E}\left[\frac{\mathbf{V}{\mathbf{E}^M}^\top \mathbf{X}}{\sqrt{n}}\left(\frac{\mathbf{V}{\mathbf{E}^M}^\top \mathbf{X}}{\sqrt{n}}\right)^\top\right] = \mathrm{tr}(\mathbf{I}_p)\mathbf{V}\boldsymbol{\Sigma}_{\boldsymbol{\varepsilon}^M}\mathbf{V}^\top, \\ &\frac{{\mathbf{E}^Y}^\top \mathbf{X}}{\sqrt{n}} \sim \mathcal{MN}(\mathbf{0}, \boldsymbol{\Sigma}_{\boldsymbol{\varepsilon}^Y}, \mathbf{I}_p), \quad \mathbb{E}\left[\frac{{\mathbf{E}^Y}^\top \mathbf{X}}{\sqrt{n}}\left(\frac{{\mathbf{E}^Y}^\top \mathbf{X}}{\sqrt{n}}\right)^\top\right] = \mathrm{tr}(\mathbf{I}_p)\boldsymbol{\Sigma}_{\boldsymbol{\varepsilon}^Y}.\end{aligned} \tag{18}$$

Plug Eq. (18) into Eq. (17) and we obtain Eq. (7). □

*Proof of Proposition 2.* The proof is similar to that of Proposition 1. The only difference is that there is no SNP-trait effect, i.e., the matrix $\mathbf{U}\mathbf{V}^\top$ for signal SNPs should be replaced by a zero matrix for noise SNPs. $\square$

*Proof of Corollary 2.* We use suffix $k$, $1 \le k \le K$, to denote the quantities associated with the $k$-th population, and suffix $\mathcal{S} \subset \{1, \dots, K\}$ to denote the quantities associated with the populations in $\mathcal{S}$. Since $\mathbf{X}_k$, $\mathbf{X}_k^0$ satisfy the assumptions in Proposition 1 and 2, then for arbitrary subset $\mathcal{S}$,

$$\begin{aligned}
\mathbf{X}_\mathcal{S}^\top \mathbf{X}_\mathcal{S} &= \sum_{k\in\mathcal{S}} \mathbf{X}_k^\top \mathbf{X}_k = \sum_{k\in\mathcal{S}} n_k \boldsymbol{I}_p = n_\mathcal{S} \boldsymbol{I}_p \\
\mathbf{X}_\mathcal{S}^{0\top} \mathbf{X}_\mathcal{S}^0 &= \sum_{k\in\mathcal{S}} \mathbf{X}_k^{0\top} \mathbf{X}_k^0 = \sum_{k\in\mathcal{S}} n_k \boldsymbol{I}_{p_0} = n_\mathcal{S} \boldsymbol{I}_{p_0}, \\
\mathbf{X}_\mathcal{S}^\top \mathbf{X}_\mathcal{S}^0 &= \sum_{k\in\mathcal{S}} \mathbf{X}_k^\top \mathbf{X}_k^0 = \sum_{k\in\mathcal{S}} \mathbf{0} = \mathbf{0}.
\end{aligned} \tag{19}$$

Immediately, we have

$$\begin{aligned}
\frac{1}{p}\mathrm{tr}\left(\mathbf{X}_\mathcal{S}^\top \mathbf{X}_\mathcal{S}\right) &= \frac{1}{p}\mathrm{tr}\left(n_\mathcal{S}\boldsymbol{I}_p\right) = n_\mathcal{S}, \\
\frac{1}{p_0}\mathrm{tr}\left(\mathbf{X}_\mathcal{S}^{0\top} \mathbf{X}_\mathcal{S}^0\right) &= \frac{1}{p_0}\mathrm{tr}\left(n_\mathcal{S}\boldsymbol{I}_{p_0}\right) = n_\mathcal{S}.
\end{aligned} \tag{20}$$

To show Corollary 2, it is sufficient to prove the case where the marginal associations can be partitioned into two parts $\boldsymbol{\beta} = [\boldsymbol{\beta}_{\mathcal{S}_1}, \boldsymbol{\beta}_{\mathcal{S}_2}]$: $\boldsymbol{\beta}_{\mathcal{S}_1}$ is based on populations $k \in \mathcal{S}_1$, and similarly for $\boldsymbol{\beta}_{\mathcal{S}_2}$. Recall model (1), (2),

$$\begin{aligned}
\boldsymbol{M} &= \boldsymbol{X}\mathbf{U} + \boldsymbol{\varepsilon}^M, \\
[\boldsymbol{Y}_{\mathcal{S}_1}, \boldsymbol{Y}_{\mathcal{S}_2}] &= \boldsymbol{M}[\mathbf{V}_{\mathcal{S}_1}^\top, \mathbf{V}_{\mathcal{S}_1}^\top] + \boldsymbol{m}[\mathbf{W}_{\mathcal{S}_1}^\top, \mathbf{W}_{\mathcal{S}_2}^\top] + [\boldsymbol{\varepsilon}^Y{}_{\mathcal{S}_1}, \boldsymbol{\varepsilon}^Y{}_{\mathcal{S}_2}],
\end{aligned}$$

then by Eq. (19), Corollary 1 is true for $\hat{\boldsymbol{\beta}}_{\mathcal{S}_1}$, $\hat{\boldsymbol{\beta}}_{\mathcal{S}_2}$, respectively,

$$\mathbb{E}\left[\hat{\boldsymbol{\beta}}_{\mathcal{S}_l}^\top \hat{\boldsymbol{\beta}}_{\mathcal{S}_l} - \frac{p}{p_0}\hat{\boldsymbol{\beta}}_{\mathcal{S}_l}^{0\top}\hat{\boldsymbol{\beta}}_{\mathcal{S}_l}^0\right] = \mathbf{V}_{\mathcal{S}_1}\mathbf{U}^\top\mathbf{U}\mathbf{V}_{\mathcal{S}_1}^\top, \quad l \in \{1, 2\}.$$

Next, we deal with the interaction term $\mathbb{E}\left[\hat{\boldsymbol{\beta}}_{\mathcal{S}_1}^\top \hat{\boldsymbol{\beta}}_{\mathcal{S}_2} - \frac{p}{p_0}\hat{\boldsymbol{\beta}}_{\mathcal{S}_1}^{0\top}\hat{\boldsymbol{\beta}}_{\mathcal{S}_2}^0\right]$. Denote the intersection of $\mathcal{S}_1$ and $\mathcal{S}_2$ by $\mathcal{S}$. Without loss of generality, we assume the first $n_\mathcal{S}$ rows of $\mathbf{X}_{\mathcal{S}_1}$, $\mathbf{X}_{\mathcal{S}_2}$ come from the populations in $\mathcal{S}$. Analogous to Eq. (15),

$$\begin{aligned}
\hat{\boldsymbol{\beta}}_{\mathcal{S}_l} &= (\mathbf{X}_{\mathcal{S}_l}^\top \mathbf{X}_{\mathcal{S}_l})^{-1}\mathbf{X}_{\mathcal{S}_l}^\top \mathbf{Y}_{\mathcal{S}_l} = \frac{\boldsymbol{X}_{\mathcal{S}_l}^\top \boldsymbol{Y}_{\mathcal{S}_l}}{n_{\mathcal{S}_l}} \\
&= \mathbf{U}\mathbf{V}_{\mathcal{S}_l}^\top + \frac{\mathbf{X}_{\mathcal{S}_l}^\top \mathbf{m}_{\mathcal{S}_l}\mathbf{W}_{\mathcal{S}_l}^\top}{n_{\mathcal{S}_l}} + \frac{\mathbf{X}_{\mathcal{S}_l}^\top \mathbf{E}^M{}_{\mathcal{S}_l}\mathbf{V}_{\mathcal{S}_l}^\top}{n_{\mathcal{S}_l}} + \frac{\mathbf{X}_{\mathcal{S}_l}^\top \mathbf{E}^Y{}_{\mathcal{S}_l}}{n_{\mathcal{S}_l}}, \quad l \in \{1, 2\}.
\end{aligned}$$

Notice that

$$\mathbb{E}\left[\frac{\mathbf{W}_{\mathcal{S}_l}\mathbf{m}_{\mathcal{S}_l}^\top \mathbf{X}_{\mathcal{S}_l}}{\sqrt{n_{\mathcal{S}_l}}}\left(\frac{\mathbf{W}_{\mathcal{S}_2}\mathbf{m}_{\mathcal{S}_2}^\top \mathbf{X}_{\mathcal{S}_2}}{\sqrt{n_{\mathcal{S}_2}}}\right)^\top\right] = \frac{\mathbf{W}_{\mathcal{S}_1}}{\sqrt{n_{\mathcal{S}_1}}}\mathbb{E}\left[\mathbf{m}_{\mathcal{S}_1}^\top \mathbf{X}_{\mathcal{S}_1}\mathbf{X}_{\mathcal{S}_2}^\top \mathbf{m}_{\mathcal{S}_2}\right]\frac{\mathbf{W}_{\mathcal{S}_2}^\top}{\sqrt{n_{\mathcal{S}_2}}},$$

and the $ij$-th element of $\mathbb{E}[\mathbf{m}_{\mathcal{S}_1}^\top \mathbf{X}_{\mathcal{S}_1} \mathbf{X}_{\mathcal{S}_2}^\top \mathbf{m}_{\mathcal{S}_2}]$ is

$$\begin{aligned}
&\mathbb{E}\left[(\mathbf{m}_{\mathcal{S}_1})_{\cdot i}^\top \mathbf{X}_{\mathcal{S}_1} \mathbf{X}_{\mathcal{S}_2}^\top (\mathbf{m}_{\mathcal{S}_2})_{\cdot j}\right] = \mathbb{E}\left[\operatorname{tr}\left(\mathbf{X}_{\mathcal{S}_1} \mathbf{X}_{\mathcal{S}_2}^\top (\mathbf{m}_{\mathcal{S}_2})_{\cdot j} (\mathbf{m}_{\mathcal{S}_1})_{\cdot i}^\top\right)\right] \\
&= \operatorname{tr}\left(\mathbf{X}_{\mathcal{S}_1} \mathbf{X}_{\mathcal{S}_2}^\top \mathbb{E}\left[(\mathbf{m}_{\mathcal{S}_2})_{\cdot j} (\mathbf{m}_{\mathcal{S}_1})_{\cdot i}^\top\right]\right) = \operatorname{tr}\left(\mathbf{X}_{\mathcal{S}_1} \mathbf{X}_{\mathcal{S}_2}^\top \begin{bmatrix} (\boldsymbol{\Sigma}_m)_{ij} I_{n_\mathcal{S}} & 0 \\ 0 & 0 \end{bmatrix}\right) \\
&= (\boldsymbol{\Sigma}_m)_{ij} \operatorname{tr}\left((\mathbf{X}_{\mathcal{S}_1})_{1:n_\mathcal{S}\cdot} (\mathbf{X}_{\mathcal{S}_2})_{1:n_\mathcal{S}\cdot}^\top\right) = (\boldsymbol{\Sigma}_m)_{ij} \operatorname{tr}\left(\mathbf{X}_\mathcal{S} \mathbf{X}_\mathcal{S}^\top\right),
\end{aligned}$$

then

$$\begin{aligned}
\mathbb{E}\left[\frac{\mathbf{W}_{\mathcal{S}_1} \mathbf{m}_{\mathcal{S}_1}^\top \mathbf{X}_{\mathcal{S}_1}}{\sqrt{n_{\mathcal{S}_1}}} \left(\frac{\mathbf{W}_{\mathcal{S}_2} \mathbf{m}_{\mathcal{S}_2}^\top \mathbf{X}_{\mathcal{S}_2}}{\sqrt{n_{\mathcal{S}_2}}}\right)^\top\right] &= \frac{n_\mathcal{S}}{\sqrt{n_{\mathcal{S}_1} n_{\mathcal{S}_2}}} \operatorname{tr}\left(\mathbf{X}_\mathcal{S} \mathbf{X}_\mathcal{S}^\top\right) \mathbf{W}_{\mathcal{S}_1} \boldsymbol{\Sigma}_m \mathbf{W}_{\mathcal{S}_2}^\top, \\
\mathbb{E}\left[\frac{\mathbf{V}_{\mathcal{S}_1} {\mathbf{E}^M}_{\mathcal{S}_1}^\top \mathbf{X}_{\mathcal{S}_1}}{\sqrt{n_{\mathcal{S}_1}}} \left(\frac{\mathbf{V}_{\mathcal{S}_2} {\mathbf{E}^M}_{\mathcal{S}_2}^\top \mathbf{X}_{\mathcal{S}_2}}{\sqrt{n_{\mathcal{S}_2}}}\right)^\top\right] &= \frac{n_\mathcal{S}}{\sqrt{n_{\mathcal{S}_1} n_{\mathcal{S}_2}}} \operatorname{tr}\left(\mathbf{X}_\mathcal{S} \mathbf{X}_\mathcal{S}^\top\right) \mathbf{V}_{\mathcal{S}_1} \boldsymbol{\Sigma}_{\boldsymbol{\varepsilon}^M} \mathbf{V}_{\mathcal{S}_2}^\top, \qquad (21) \\
\mathbb{E}\left[\frac{{\mathbf{E}^Y}_{\mathcal{S}_1}^\top \mathbf{X}_{\mathcal{S}_1}}{\sqrt{n_{\mathcal{S}_1}}} \left(\frac{{\mathbf{E}^Y}_{\mathcal{S}_2}^\top \mathbf{X}_{\mathcal{S}_2}}{\sqrt{n_{\mathcal{S}_{\mathcal{S}_2}}}}\right)^\top\right] &= \frac{n_\mathcal{S}}{\sqrt{n_{\mathcal{S}_1} n_{\mathcal{S}_2}}} \operatorname{tr}\left(\mathbf{X}_\mathcal{S} \mathbf{X}_\mathcal{S}^\top\right) \mathsf{Cov}\left({\boldsymbol{\varepsilon}^Y}_{\mathcal{S}_1}, {\boldsymbol{\varepsilon}^Y}_{\mathcal{S}_2}\right).
\end{aligned}$$

Eq. (21) is valid if we replace signal SNP matrices $\mathbf{X}_{\mathcal{S}_1}$, $\mathbf{X}_{\mathcal{S}_2}$ by noise SNP matrices $\mathbf{X}_{\mathcal{S}_1}^0$, $\mathbf{X}_{\mathcal{S}_2}^0$. Therefore, by Eq. (20) and (21),

$$\mathbb{E}\left[\hat{\boldsymbol{\beta}}_{\mathcal{S}_1}^\top \hat{\boldsymbol{\beta}}_{\mathcal{S}_2} - \frac{p}{p_0} \hat{\boldsymbol{\beta}}_{\mathcal{S}_1}^{0\top} \hat{\boldsymbol{\beta}}_{\mathcal{S}_2}^0\right] = \mathbf{V}_{\mathcal{S}_1} \mathbf{U}^\top \mathbf{U} \mathbf{V}_{\mathcal{S}_2}^\top.$$

Finally, merge the results of diagonal and off-diagonal sub-matrices,

$$\begin{aligned}
\mathbb{E}\left[\hat{\boldsymbol{\beta}}^\top \hat{\boldsymbol{\beta}} - \frac{p}{p_0} \hat{\boldsymbol{\beta}}^{0\top} \hat{\boldsymbol{\beta}}^0\right] &= \begin{bmatrix} \hat{\boldsymbol{\beta}}_{\mathcal{S}_1}^\top \hat{\boldsymbol{\beta}}_{\mathcal{S}_1} - \frac{p}{p_0} \hat{\boldsymbol{\beta}}_{\mathcal{S}_1}^{0\top} \hat{\boldsymbol{\beta}}_{\mathcal{S}_1}^0 & \hat{\boldsymbol{\beta}}_{\mathcal{S}_1}^\top \hat{\boldsymbol{\beta}}_{\mathcal{S}_2} - \frac{p}{p_0} \hat{\boldsymbol{\beta}}_{\mathcal{S}_1}^{0\top} \hat{\boldsymbol{\beta}}_{\mathcal{S}_2}^0 \\ \hat{\boldsymbol{\beta}}_{\mathcal{S}_2}^\top \hat{\boldsymbol{\beta}}_{\mathcal{S}_1} - \frac{p}{p_0} \hat{\boldsymbol{\beta}}_{\mathcal{S}_2}^{0\top} \hat{\boldsymbol{\beta}}_{\mathcal{S}_1}^0 & \hat{\boldsymbol{\beta}}_{\mathcal{S}_2}^\top \hat{\boldsymbol{\beta}}_{\mathcal{S}_2} - \frac{p}{p_0} \hat{\boldsymbol{\beta}}_{\mathcal{S}_2}^{0\top} \hat{\boldsymbol{\beta}}_{\mathcal{S}_2}^0 \end{bmatrix} \\
&= \begin{bmatrix} \mathbf{V}_{\mathcal{S}_1} \mathbf{U}^\top \mathbf{U} \mathbf{V}_{\mathcal{S}_1}^\top & \mathbf{V}_{\mathcal{S}_1} \mathbf{U}^\top \mathbf{U} \mathbf{V}_{\mathcal{S}_2}^\top \\ \mathbf{V}_{\mathcal{S}_2} \mathbf{U}^\top \mathbf{U} \mathbf{V}_{\mathcal{S}_1}^\top & \mathbf{V}_{\mathcal{S}_2} \mathbf{U}^\top \mathbf{U} \mathbf{V}_{\mathcal{S}_2}^\top \end{bmatrix} = \begin{bmatrix} \mathbf{V}_{\mathcal{S}_1} \\ \mathbf{V}_{\mathcal{S}_2} \end{bmatrix} \mathbf{U}^\top \mathbf{U} \begin{bmatrix} \mathbf{V}_{\mathcal{S}_1} \\ \mathbf{V}_{\mathcal{S}_2} \end{bmatrix}^\top = \mathbf{V} \mathbf{U}^\top \mathbf{U} \mathbf{V}^\top,
\end{aligned}$$

and we finish the proof. □

### 5 Data description

#### 5.1 Metabolomics Data

The main metabolomics dataset [KDW+16] contains 123 metabolites and around $1.3 \times 10^7$ SNPs. Figure 5 illustrates the preprocessing procedure of the metabolomics data. We describe each step in detail. Table 2 summarizes the 105 metabolites after preprocessing.

a Select index SNPs. We use an external dataset [DHL+17] of 72 metabolites to select independent representative SNPs or index SNPs. The external dataset contains a large proportion of the lipoprotein measures in the main dataset.

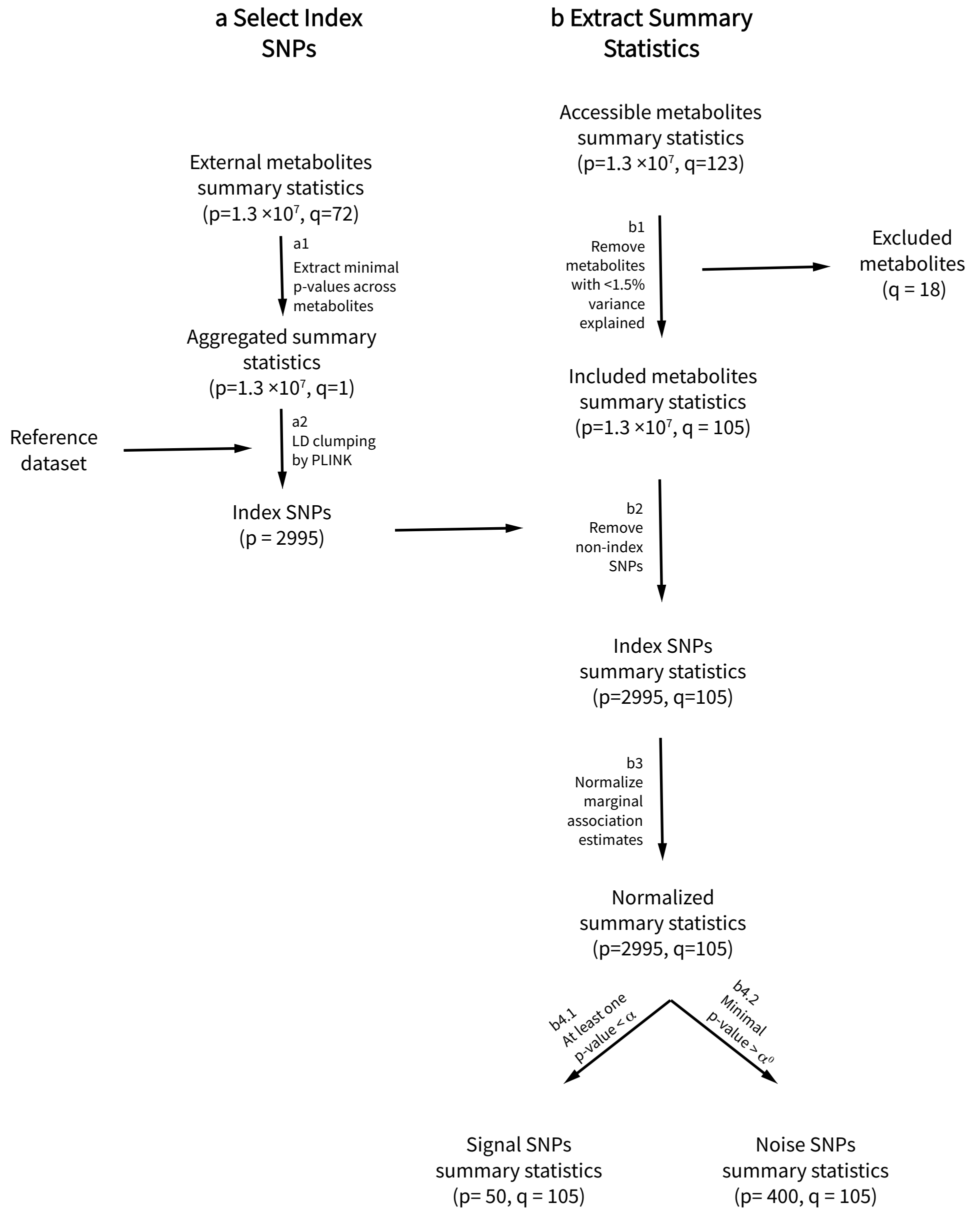

Figure 5: *Preprocessing of metabolomics data.*

a1 *Extract minimal p-values across traits.* In the external dataset, for each SNP, we compute the minimal p-value across all 72 marginal associations. We form a dataset of SNPs and minimal p-values and call it the aggregated summary statistics dataset.

a2 *LD clumping by PLINK [PNTB+07].* We use the PLINK software to form independent SNP groups and select index SNPs according to the minimal p-values. PLINK requires a reference dataset of empirical estimates of linkage disequilibrium between SNPs and we use 1000 genome European reference panel dataset [LAE+21, ELA+20]. There are four main parameters determining the level of clumping: the significance threshold for index SNPs, the secondary significance threshold for clumped SNPs, LD threshold for clumping, and physical distance threshold for clumping. We remark that our goal is to form independent SNP groups and do not require index SNPs to be significant, thus we set significance thresholds at 1, i.e., no thresholding, and choose a small LD threshold 0.001 and a large physical distance 10000. The PLINK software outputs 2995 SNP clumps, and we regard the associated 2995 index SNPs as independent and use them for the downstream analysis.

b Extract and clean summary statistics for signal and noise SNPs. We work with the main metabolomics dataset below.

b1 *Remove metabolites with low variance explained.* We discard traits with less than 1.5% variance explained by genes using the proportion of variance explained in the Supplementary Table 1 of [KDW+16]. There are 18 out of 123 metabolites excluded, accounting for around 15% of the traits. Most of the removed metabolites are small molecules or protein measures.

b2 *Remove non-index SNPs.* We only keep the summary statistics corresponding to the included 105 metabolites from step b1 and the 2995 index SNPs from step a2.

b3 *Normalize marginal association estimates.* For each metabolite, we divide all the relevant marginal association estimates by the median of standard deviations over 2995 index SNPs. The normalized marginal associations are invariant to the response (metabolites measurements) scaling.

b4.1 *Select signal SNPs.* We call a SNP signal SNP if there is at least one marginal association test significant at level $\alpha = 10^{-6}/105$. Here we use $10^{-6}$ as the significance level for one trait and apply Bonferroni correction. Finally, we obtain 50 independent signal SNPs.

b4.2 *Select noise SNPs.* We call a SNP noise SNP if all the marginal association tests are insignificant at level $\alpha^0$—the 5% quantile of 105 independent uniform $[0, 1]$ random variables. The rationale is that, if an index SNP does not influence any of the 105 traits, the associated p-values should be uniformly distributed on $[0, 1]$. The above $\alpha^0$ guarantees at most 5% noise SNPs will be regarded as signal SNPs when the p-values are approximately independent. We end up with 400 independent noise SNPs.

#### 5.2 UK Biobank Data

We use the GWAS summary statistics from UK biobank data: UKBB GWAS Imputed v3 generated by Neale Lab (http://www.nealelab.is/uk-biobank). The dataset is based on $3.6 \times 10^5$

| Abbreviation | | Description |
|---|---|---|
| | | *Lipoprotein measures* |
| M.LDL.P | HDL.C | In lipoprotein abbreviations, |
| M.VLDL.CE | HDL.D | First is given the size category if applicable: |
| M.VLDL.C | IDL.C | XL=extra large, |
| M.VLDL.FC | IDL.FC | L=large, |
| M.VLDL.L | IDL.L | M=medium, |
| M.VLDL.PL | IDL.PL | S=small |
| M.VLDL.P | IDL.P | XS=extra small. |
| M.VLDL.TG | IDL.TG | Second the lipoprotein particle is given: |
| S.HDL.L | LDL.C | VLDL=very-low-density lipoprotein particle, |
| S.HDL.P | LDL.D | IDL= intermediate-density lipoprotein particle, |
| S.HDL.TG | L.HDL.CE | LDL=low-density lipoprotein particles, |
| S.LDL.C | L.HDL.C | HDL=high-density lipoprotein particle. |
| S.LDL.L | L.HDL.FC | Third the lipid measure of the particle: |
| S.LDL.P | L.HDL.L | C=total cholesterol, |
| S.VLDL.C | L.HDL.PL | D=the mean diameter of the particle, |
| S.VLDL.FC | L.HDL.P | FC=free cholesterol, |
| S.VLDL.L | L.LDL.CE | L=total lipids, |
| S.VLDL.PL | L.LDL.C | P=particle concentration, |
| S.VLDL.P | L.LDL.FC | PL=phospholipids, |
| S.VLDL.TG | L.LDL.L | TG= triglycerides. |
| VLDL.D | L.LDL.PL | |
| XL.HDL.CE | L.LDL.P | |
| XL.HDL.C | L.VLDL.CE | |
| XL.HDL.FC | L.VLDL.C | |
| XL.HDL.L | L.VLDL.FC | |
| XL.HDL.PL | L.VLDL.L | |
| XL.HDL.P | L.VLDL.PL | |
| XL.HDL.TG | L.VLDL.P | |
| XL.VLDL.L | L.VLDL.TG | |
| XL.VLDL.PL | M.HDL.CE | |
| XL.VLDL.P | M.HDL.C | |
| XL.VLDL.TG | M.HDL.FC | |
| XS.VLDL.L | M.HDL.L | |
| XS.VLDL.PL | M.HDL.PL | |
| XS.VLDL.P | M.HDL.P | |
| XS.VLDL.TG | M.LDL.CE | |
| XXL.VLDL.L | M.LDL.C | |
| XXL.VLDL.PL | M.LDL.L | |
| XXL.VLDL.TG | M.LDL.PL | |

Table 2: *Table of metabolites' abbreviations and descriptions.*

| Abbreviation | Description |
|---|---|
| | *Lipid measures* |
| Bis.DB.ratio | Ratio of bis-allylic bonds to double bonds in lipids |
| Bis.FA.ratio | Ratio of bis-allylic bonds to total fatty acids in lipids |
| CH2.DB.ratio | CH2 groups to double bonds ratio |
| CH2.in.FA | CH2 groups in fatty acids |
| DB.in.FA | Double bonds in fatty acids |
| DHA | docosahexaenoic acid |
| Est.C | Esterified cholesterol |
| FALen | Fatty acid length |
| FAw3 | Omega-3 fatty acids |
| FAw6 | Omega-6 fatty acids |
| FAw79S | Omega-7 and -9 and saturated fatty acids |
| FreeC | Free cholesterol |
| LA | Linoleic acid |
| MUFA | Mono-unsaturated fatty acids |
| otPUFA | Other polyunsaturated fatty acids |
| PC | Phosphatidylcholine and other cholines |
| Serum.C | Serum total cholesterol |
| Serum.TG | Serum total triglycerides |
| SM | Sphingomyelins |
| Tot.FA | Total fatty acids |
| Tot.PG | Total phosphoglycerides |
| | *Small molecules and protein measures* |
| ApoA1 | Apolipoprotein A1 |
| APOB | Apolipoprotein B |
| Gln | Glutamine |
| Glol | Glycerol |
| Gly | Glycine |
| Gp | Glycoprotein acetyls, mainly a1-acid glycoprotein |

*Table of metabolites' abbreviations and descriptions (continued).*

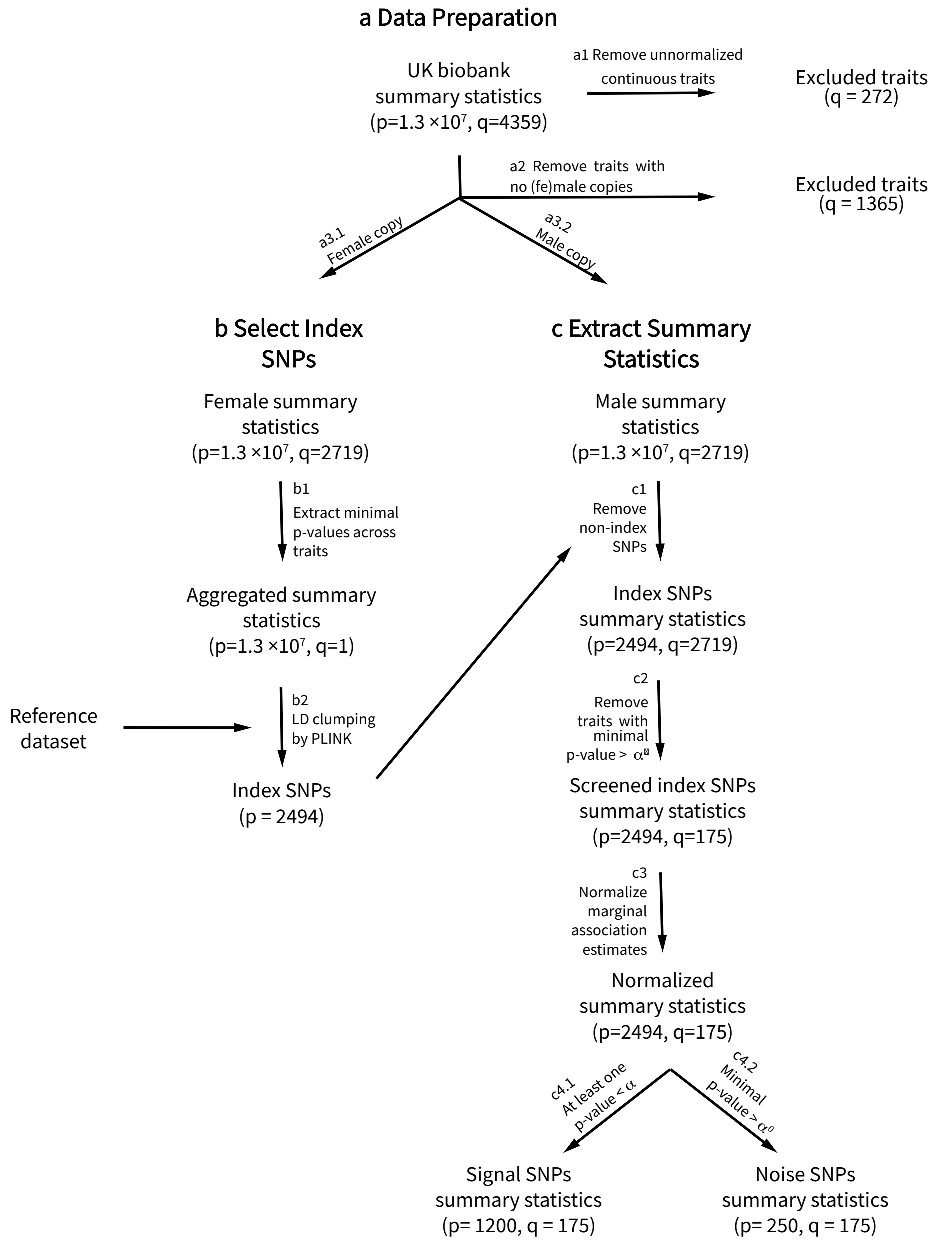

Figure 6: *Preprocessing of UK biobank data.*

samples of white-British ancestry and $1.3 \times 10^7$ QC-passing SNPs. Figure 5 illustrates our preprocessing procedure of the UK biobank data. Below we describe each step in detail. Table 3 summarizes the 175 traits after preprocessing. Briefly, the resulting traits are a combination of lab measurements, diagnoses and medication, and a small number of living habits.

a Data preparation.

a1 For each continuous variable, two versions are available: an untransformed version ("continuous_raw") and a version where values are inverse rank normalized ("continuous_irnt"). We adopt the transformed version and remove 272 unnormalized counterparts.

a2 The total sample of $3.6 \times 10^5$ individuals contains $1.9 \times 10^5$ female subjects and $1.7 \times 10^5$ male subjects. Over two-thirds of the traits come with three summary statistics corresponding to the full sample, the female subset, and the male subset. We keep the traits with both female and male summary statistics and remove 1365 traits with no female or male summary statistics. The female and male copies will be of different use in the downstream analysis.

a3.1 We use the female summary statistics for index SNP selection.

a3.2 We use the male summary statistics to construct our major summary statistics matrix.

b Select index SNPs. We use the female summary statistics to select independent index SNPs.

b1 *Extract minimal p-values across traits.* In the female dataset, for each SNP, we compute the minimal p-value across all 2719 statistical tests of marginal associations. We form a dataset of SNPs and minimal p-values and call it the aggregated summary statistics dataset.

b2 *LD clumping by PLINK [PNTB+07].* We use the PLINK software to form independent SNP groups and select index SNPs according to the minimal p-values. Details are discussed in Section 5.1. The PLINK software outputs 2494 SNP clumps and the associated index SNPs.

c Extract and clean summary statistics for signal and noise SNPs. We work with the male summary statistics below.

c1 *Remove non-index SNPs.* We only keep the summary statistics corresponding to the 2494 index SNPs from step b2.

c2 *Remove traits with no significant index SNPs.* We remove traits with no index SNP significant at level $\alpha^P = 5 \times 10^{-8}/2494$. Many living habits are excluded. We are left with 175 traits.

c3 *Normalize marginal association estimates.* For each trait, we divide all the marginal association estimates by the median of standard deviations over 2494 index SNPs.

c4.1 *Select signal SNPs.* We select signal SNPs with at least one marginal association test significant at level $\alpha = 10^{-6}/175$. We obtain 1200 independent signal SNPs. More details are discussed in Section 5.1.

c4.2 *Select noise SNPs.* We select noise SNPS if all the marginal association tests are insignificant at level $\alpha^0$—5% quantile of 175 independent uniform $[0, 1]$ random variables. We obtain 250 independent noise SNPs. More details are discussed in Section 5.1.

| Trait code | Description |
|---|---|
| 102_irnt | Pulse rate, automated reading |
| 1249 | Past tobacco smoking |
| 1558 | Alcohol intake frequency |
| 1717 | Skin colour |
| 1727 | Ease of skin tanning |
| 1737 | Childhood sunburn occasions |
| 1747_1 | Hair colour (natural, before greying): Blonde |
| 1747_2 | Hair colour (natural, before greying): Red |
| 1747_3 | Hair colour (natural, before greying): Light brown |
| 1747_4 | Hair colour (natural, before greying): Dark brown |
| 1747_5 | Hair colour (natural, before greying): Black |
| 1747_6 | Hair colour (natural, before greying): Other |
| 1757 | Facial aging |
| 20002_1065 | Non-cancer illness code, self-reported: hypertension |
| 20002_1093 | Non-cancer illness code, self-reported: pulmonary embolism +/- dvt |
| 20002_1094 | Non-cancer illness code, self-reported: deep venous thrombosis (dvt) |
| 20002_1111 | Non-cancer illness code, self-reported: asthma |
| 20002_1220 | Non-cancer illness code, self-reported: diabetes |
| 20002_1223 | Non-cancer illness code, self-reported: type 2 diabetes |
| 20002_1226 | Non-cancer illness code, self-reported: hypothyroidism/myxoedema |
| 20002_1445 | Non-cancer illness code, self-reported: clotting disorder/excessive bleeding |
| 20002_1451 | Non-cancer illness code, self-reported: hereditary/genetic haematological disorder |
| 20002_1471 | Non-cancer illness code, self-reported: atrial fibrillation |
| 20002_1473 | Non-cancer illness code, self-reported: high cholesterol |
| 20003_1140861924 | Treatment/medication code: bezafibrate |
| 20003_1140861954 | Treatment/medication code: fenofibrate |
| 20003_1140861958 | Treatment/medication code: simvastatin |
| 20003_1140861998 | Treatment/medication code: ventolin 100micrograms inhaler |
| 20003_1140874744 | Treatment/medication code: gliclazide |
| 20003_1140883066 | Treatment/medication code: insulin product |
| 20003_1140884600 | Treatment/medication code: metformin |
| 20003_1140888266 | Treatment/medication code: warfarin |
| 20003_1140888570 | Treatment/medication code: flecainide |
| 20003_1141146138 | Treatment/medication code: lipitor 10mg tablet |
| 20003_1141146234 | Treatment/medication code: atorvastatin |
| 20003_1141171646 | Treatment/medication code: pioglitazone |
| 20003_1141191044 | Treatment/medication code: levothyroxine sodium |
| 20003_1141192736 | Treatment/medication code: ezetimibe |
| 20003_2038459814 | Treatment/medication code: digoxin |
| 20015_irnt | Sitting height |

Table 3: *Table of UK biobank trait codes and descriptions.*

| Trait code | Description |
| --- | --- |
| 20022_irnt | Birth weight |
| 20116_0 | Smoking status: Never |
| 20116_1 | Smoking status: Previous |
| 20150_irnt | Forced expiratory volume in 1-second (FEV1), Best measure |
| 20151_irnt | Forced vital capacity (FVC), Best measure |
| 20153_irnt | Forced expiratory volume in 1-second (FEV1), predicted |
| 20154_irnt | Forced expiratory volume in 1-second (FEV1), predicted percentage |
| 20160 | Ever smoked |
| 20403 | Amount of alcohol drunk on a typical drinking day |
| 20406 | Ever addicted to alcohol |
| 20414 | Frequency of drinking alcohol |
| 20416 | Frequency of consuming six or more units of alcohol |
| 21001_irnt | Body mass index (BMI) |
| 21002_irnt | Weight |
| 22127 | Doctor diagnosed asthma |
| 2247_0 | Hearing difficulty/problems: No |
| 2247_1 | Hearing difficulty/problems: Yes |
| 2267 | Use of sun/uv protection |
| 23098_irnt | Weight |
| 23099_irnt | Body fat percentage |
| 23100_irnt | Whole body fat mass |
| 23101_irnt | Whole body fat-free mass |
| 23102_irnt | Whole body water mass |
| 23104_irnt | Body mass index (BMI) |
| 23105_irnt | Basal metabolic rate |
| 23111_irnt | Leg fat percentage (right) |
| 23112_irnt | Leg fat mass (right) |
| 23113_irnt | Leg fat-free mass (right) |
| 23114_irnt | Leg predicted mass (right) |
| 23115_irnt | Leg fat percentage (left) |
| 23116_irnt | Leg fat mass (left) |
| 23117_irnt | Leg fat-free mass (left) |
| 23118_irnt | Leg predicted mass (left) |
| 23119_irnt | Arm fat percentage (right) |
| 23120_irnt | Arm fat mass (right) |
| 23121_irnt | Arm fat-free mass (right) |
| 23122_irnt | Arm predicted mass (right) |
| 23123_irnt | Arm fat percentage (left) |
| 23124_irnt | Arm fat mass (left) |
| 23125_irnt | Arm fat-free mass (left) |

*Table of UK biobank trait codes and descriptions (continued).*

| Trait code | Description |
|---|---|
| 23126_irnt | Arm predicted mass (left) |
| 23127_irnt | Trunk fat percentage |
| 23128_irnt | Trunk fat mass |
| 23129_irnt | Trunk fat-free mass |
| 23130_irnt | Trunk predicted mass |
| 2316 | Wheeze or whistling in the chest in last year |
| 2443 | Diabetes diagnosed by doctor |
| 2887 | Number of cigarettes previously smoked daily |
| 2986 | Started insulin within one year diagnosis of diabetes |
| 30000_irnt | White blood cell (leukocyte) count |
| 30010_irnt | Red blood cell (erythrocyte) count |
| 30020_irnt | Haemoglobin concentration |
| 30030_irnt | Haematocrit percentage |
| 30040_irnt | Mean corpuscular volume |
| 30050_irnt | Mean corpuscular haemoglobin |
| 30060_irnt | Mean corpuscular haemoglobin concentration |
| 30070_irnt | Red blood cell (erythrocyte) distribution width |
| 30080_irnt | Platelet count |
| 30090_irnt | Platelet crit |
| 30100_irnt | Mean platelet (thrombocyte) volume |
| 30110_irnt | Platelet distribution width |
| 30120_irnt | Lymphocyte count |
| 30130_irnt | Monocyte count |
| 30140_irnt | Neutrophill count |
| 30150 | Eosinophill count |
| 30160 | Basophill count |
| 30180_irnt | Lymphocyte percentage |
| 30190_irnt | Monocyte percentage |
| 30200_irnt | Neutrophill percentage |
| 30210_irnt | Eosinophill percentage |
| 30220_irnt | Basophill percentage |
| 30240_irnt | Reticulocyte percentage |
| 30250_irnt | Reticulocyte count |
| 30260_irnt | Mean reticulocyte volume |
| 30270_irnt | Mean sphered cell volume |
| 30280_irnt | Immature reticulocyte fraction |
| 30290_irnt | High light scatter reticulocyte percentage |
| 30300_irnt | High light scatter reticulocyte count |
| 3062_irnt | Forced vital capacity (FVC) |
| 3063_irnt | Forced expiratory volume in 1-second (FEV1) |

*Table of UK biobank trait codes and descriptions (continued).*

| Trait code | Description |
|---|---|
| 3064_irnt | Peak expiratory flow (PEF) |
| 3144_irnt | Heel Broadband ultrasound attenuation, direct entry |
| 3147_irnt | Heel quantitative ultrasound index (QUI), direct entry |
| 3148_irnt | Heel bone mineral density (BMD) |
| 3476 | Difficulty not smoking for 1 day |
| 4079_irnt | Diastolic blood pressure, automated reading |
| 4080_irnt | Systolic blood pressure, automated reading |
| 4101_irnt | Heel broadband ultrasound attenuation (left) |
| 4104_irnt | Heel quantitative ultrasound index (QUI), direct entry (left) |
| 4105_irnt | Heel bone mineral density (BMD) (left) |
| 4106_irnt | Heel bone mineral density (BMD) T-score, automated (left) |
| 4119_irnt | Ankle spacing width (right) |
| 4120_irnt | Heel broadband ultrasound attenuation (right) |
| 4123_irnt | Heel quantitative ultrasound index (QUI), direct entry (right) |
| 4124_irnt | Heel bone mineral density (BMD) (right) |
| 4125_irnt | Heel bone mineral density (BMD) T-score, automated (right) |
| 4194_irnt | Pulse rate |
| 48_irnt | Waist circumference |
| 49_irnt | Hip circumference |
| 50_irnt | Standing height |
| 5983_irnt | ECG, heart rate |
| 6033_irnt | Maximum heart rate during fitness test |
| 6148_1 | Eye problems/disorders: Diabetes related eye disease |
| 6150_4 | Vascular/heart problems diagnosed by doctor: High blood pressure |
| 6152_5 | Blood clot, DVT, bronchitis, emphysema, asthma, rhinitis, eczema, allergy diagnosed by doctor: Blood clot in the leg (DVT) |
| 6152_7 | Blood clot, DVT, bronchitis, emphysema, asthma, rhinitis, eczema, allergy diagnosed by doctor: Blood clot in the lung |
| 6152_8 | Blood clot, DVT, bronchitis, emphysema, asthma, rhinitis, eczema, allergy diagnosed by doctor: Asthma |
| 6154_3 | Medication for pain relief, constipation, heartburn: Paracetamol |
| 78_irnt | Heel bone mineral density (BMD) T-score, automated |
| C_OTHER_SKIN | Other malignant neoplasms of skin |
| C_SKIN | NA |
| C3_OTHER_SKIN | Other malignant neoplasms of skin |
| C3_SKIN | Malignant neoplasm of skin |
| C44 | Diagnoses - main ICD10: C44 Other malignant neoplasms of skin |
| CARDIAC _ARRHYTM | Cardiac arrhytmias, COPD co-morbidities |
| E83 | Diagnoses - main ICD10: E83 Disorders of mineral metabolism |
| I21 | Diagnoses - main ICD10: I21 Acute myocardial infarction |
| I25 | Diagnoses - main ICD10: I25 Chronic ischaemic heart disease |
| I26 | Diagnoses - main ICD10: I26 Pulmonary embolism |
| I48 | Diagnoses - main ICD10: I48 Atrial fibrillation and flutter |

*Table of UK biobank trait codes and descriptions (continued).*

| Trait code | Description |
|---|---|
| I80 | Diagnoses - main ICD10: I80 Phlebitis and thrombophlebitis |
| I83 | Diagnoses - main ICD10: I83 Varicose veins of lower extremities |
| I9_CHD | Major coronary heart disease event |
| I9_CHD_NOREV | Major coronary heart disease event excluding revascularizations |
| I9_CORATHER | Coronary atherosclerosis |
| I9_DISVEINLYMPH | Diseases of veins, lymphatic vessels and lymph nodes, not elsewhere classified |
| I9_DVTANDPULM | DVT of lower extremities and pulmonary embolism |
| I9_IHD | Ischaemic heart disease, wide definition |
| I9_MI | Myocardial infarction |
| I9_MI_STRICT | Myocardial infarction, strict |
| I9_PHLETHRO -MBDVTLOW | DVT of lower extremities |
| I9_VTE | Venous thromboembolism |
| I95 | Diagnoses - main ICD10: I95 Hypotension |
| IX_CIRCULATORY | Diseases of the circulatory system |
| OTHER_ILD_CVD _COMORB | Other ILD-related CVD-co-morbidities |

*Table of UK biobank trait codes and descriptions (continued).*

### 6 Auxiliary result

#### 6.1 Gene-trait clusters

We display details of gene-trait clusters learnt from the metabolomics data and UK biobank data. We follow the co-appearance graph clustering with bagging (Algorithm 2) and consider three clustering methods: UMAP embedding with k-means, t-SNE embedding with k-means, and spectral clustering.

For the metabolomics data, Figures 7 and 8 display the 2-dimensional embeddings from UMAP and t-SNE, respectively. Genes and traits are colored according to the clustering memberships. We connect a gene and a trait if the associated marginal association estimate is the 100 most significant. In Figure 9, we demonstrate the spectral clustering results using the co-appearance heatmap grouped by the clustering membership. Tables 4, 5, and 6 list the detailed clustering results. For each SNP, we obtain relevant information using R-package haploR [WK12]. We input reference SNP cluster IDs (rsIDs) to haploR and receive associated candidate genes and organisms where the SNP is significantly associated with the expression of the candidate genes. We remark that for each variant, haploR calculates the proximity of the variant to genes by either annotation or the orientation relative to the nearest end of the genes and outputs the gene with the highest proximity as the candidate gene.

Similarly for the UK biobank data, Figures 10, 11, and 12 display the clustering results from UMAP, t-SNE, and spectral clustering, respectively. We list the detailed clustering results in separate CSV files.

#### 6.2 Stability results

We numerically demonstrate the robustness of bootstrap aggregation to trait collection discussed in Section 2.2. In practice, various datasets with moderately different collections of traits may address the same biological question, and it is desirable to arrive at consistent conclusions despite the difference in trait collection. In this section, given a dataset, we artificially create different versions of the dataset through subsampling traits without replacement and evaluate the congruence of clustering results across subsamples.

We discuss the procedure applied to the metabolomics data in detail and the UK biobank data is handled similarly. We start from the marginal association estimate matrix of index SNPs with a total of 105 traits. In each trial, we fix the index SNPs and subsample 50 traits without replacement. We then apply the following analysis to the submatrix:

- One-shot baseline (Algorithm 1).
- Co-appearance graph clustering without bootstrap aggregation. We use estimators $\hat{\mathbf{U}}$, $\hat{\mathbf{V}}$ from the one-shot baseline to construct co-appearance graphs. We then apply three clustering methods: UMAP with k-means, t-SNE with k-means, and spectral clustering to the co-appearance graphs.
- Co-appearance graph clustering with bootstrap aggregation (Algorithm 2). We use the same clustering methods as the co-appearance frequency graph clustering without bootstrap aggregation.

To make comparisons fair, we fix all shared hyper-parameters at the same values.

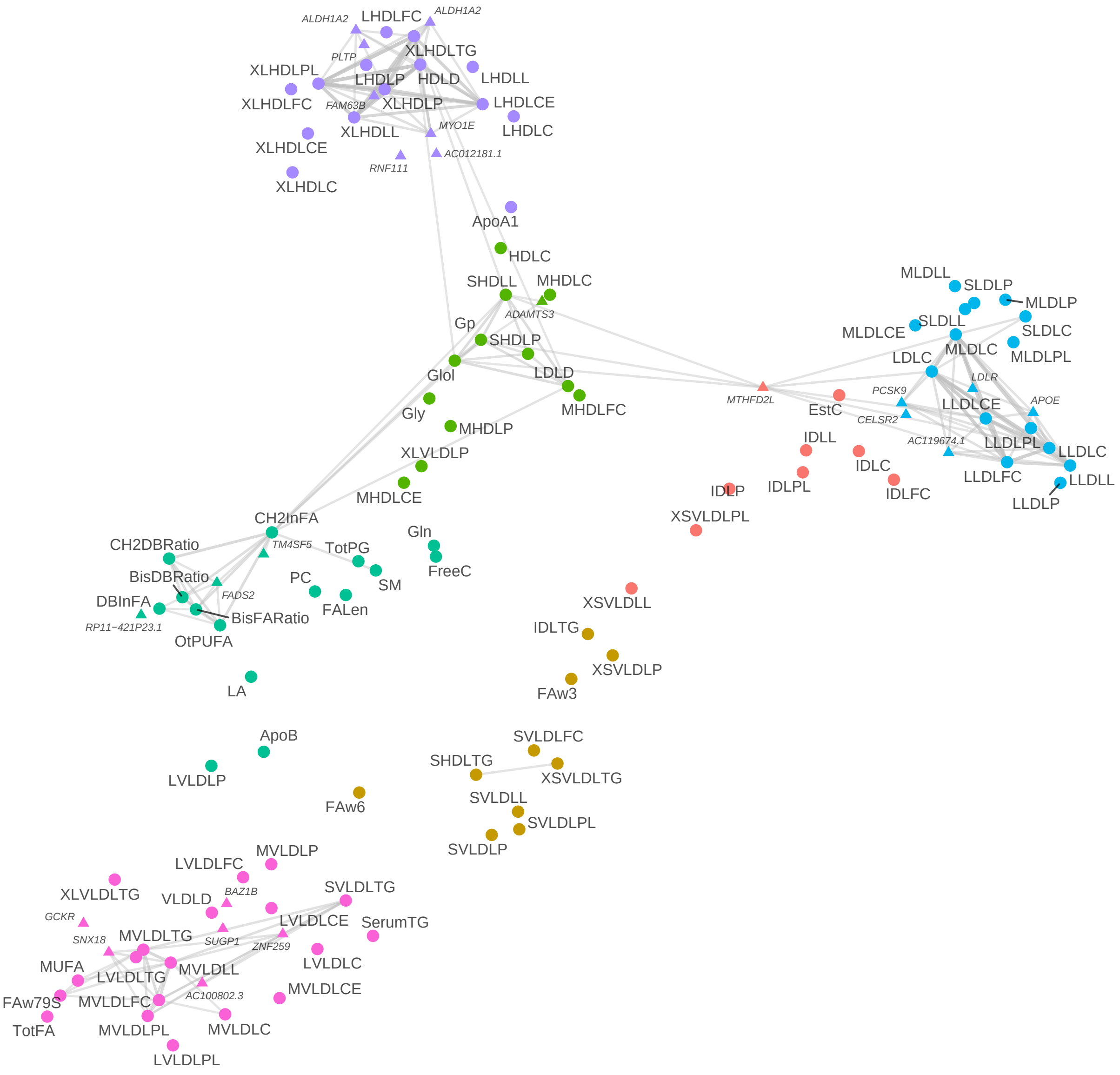

Figure 7: *Visualization of gene-trait clusters based on the metabolomics data using UMAP and kmeans. Circle points represent traits and triangle points represent candidate genes of signal index SNPs. A gene and a trait is connected by a grey line if the associated marginal association estimate is among the top* 100 *significant.*

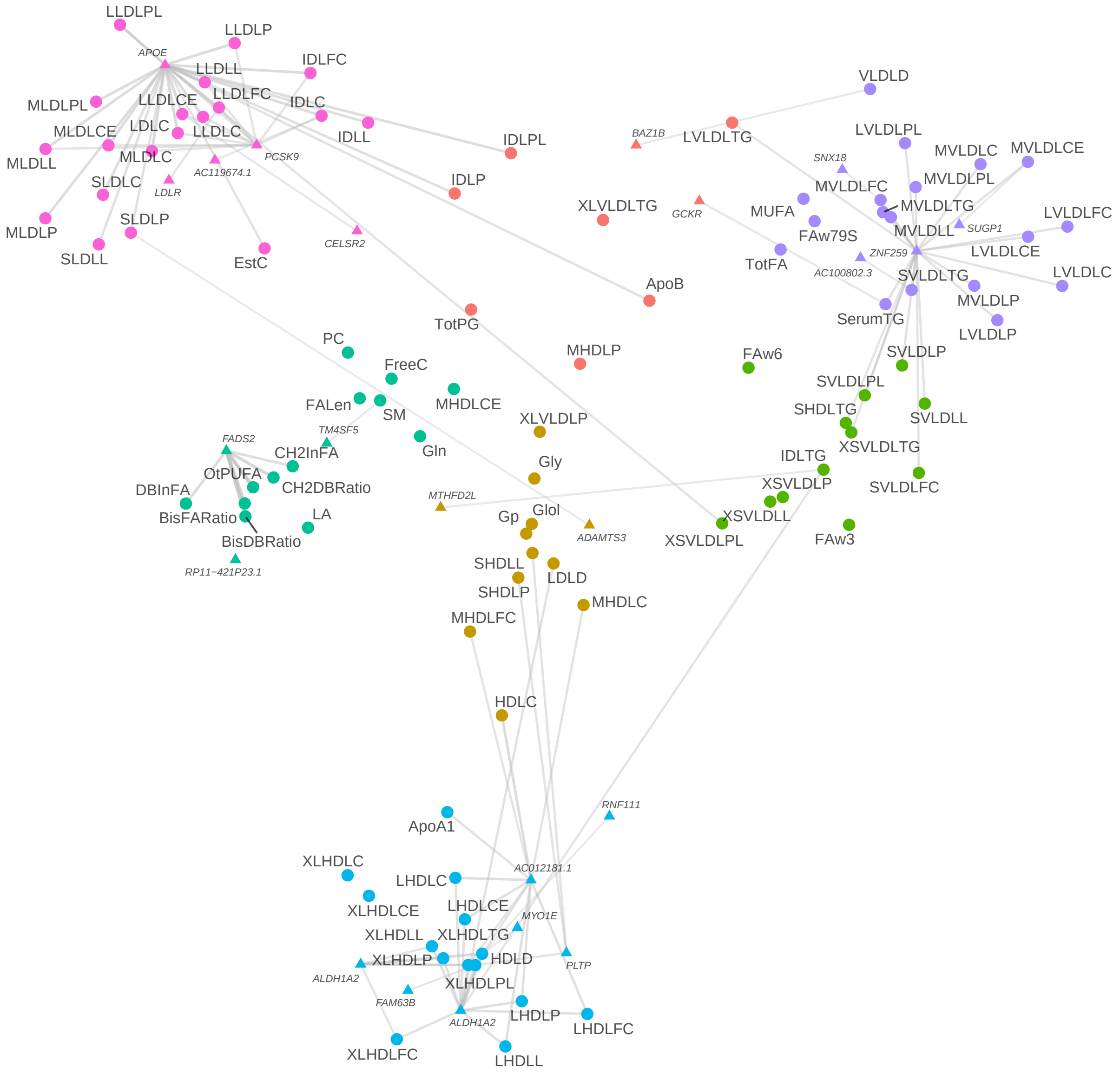

Figure 8: *Visualization of gene-trait clusters based on the metabolomics data using t-SNE and kmeans. More details can be found in the caption of Figure 7.*

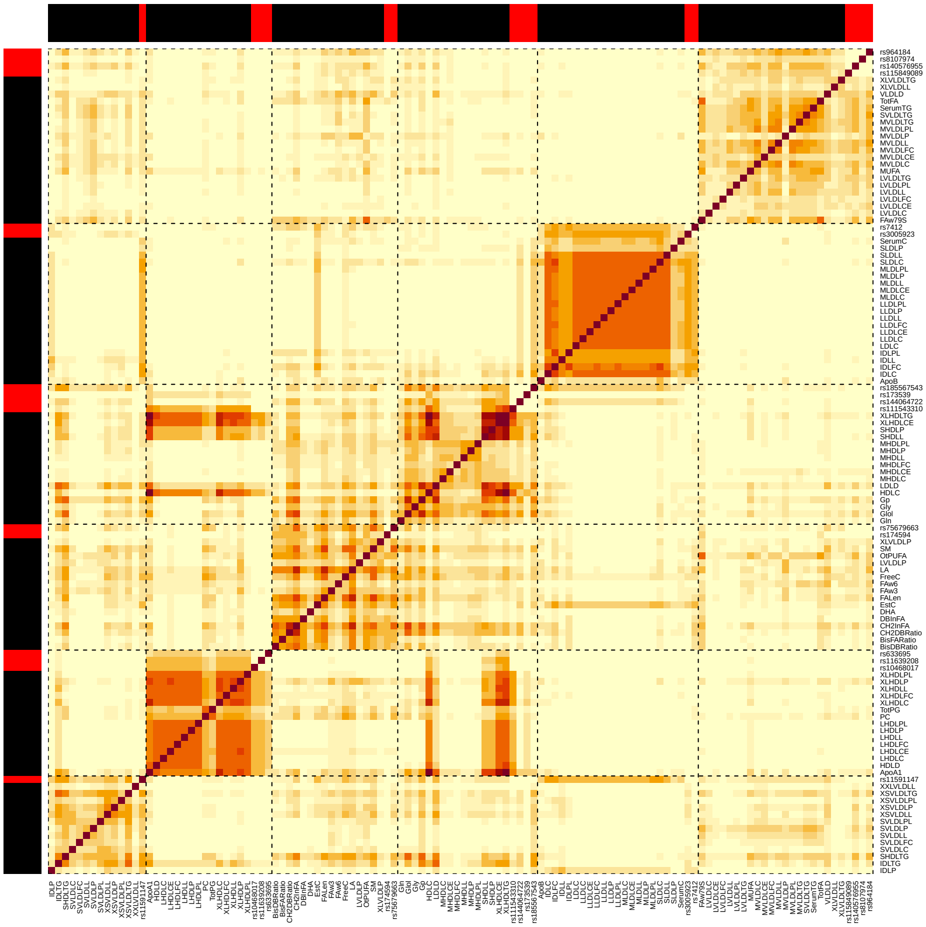

Figure 9: *Co-appearance heatmap based on the spectral clustering results on the metabolomics data. The dashed lines separate different clusters. The horizontal and vertical side bars use black to denote traits and red to denote SNPs.*

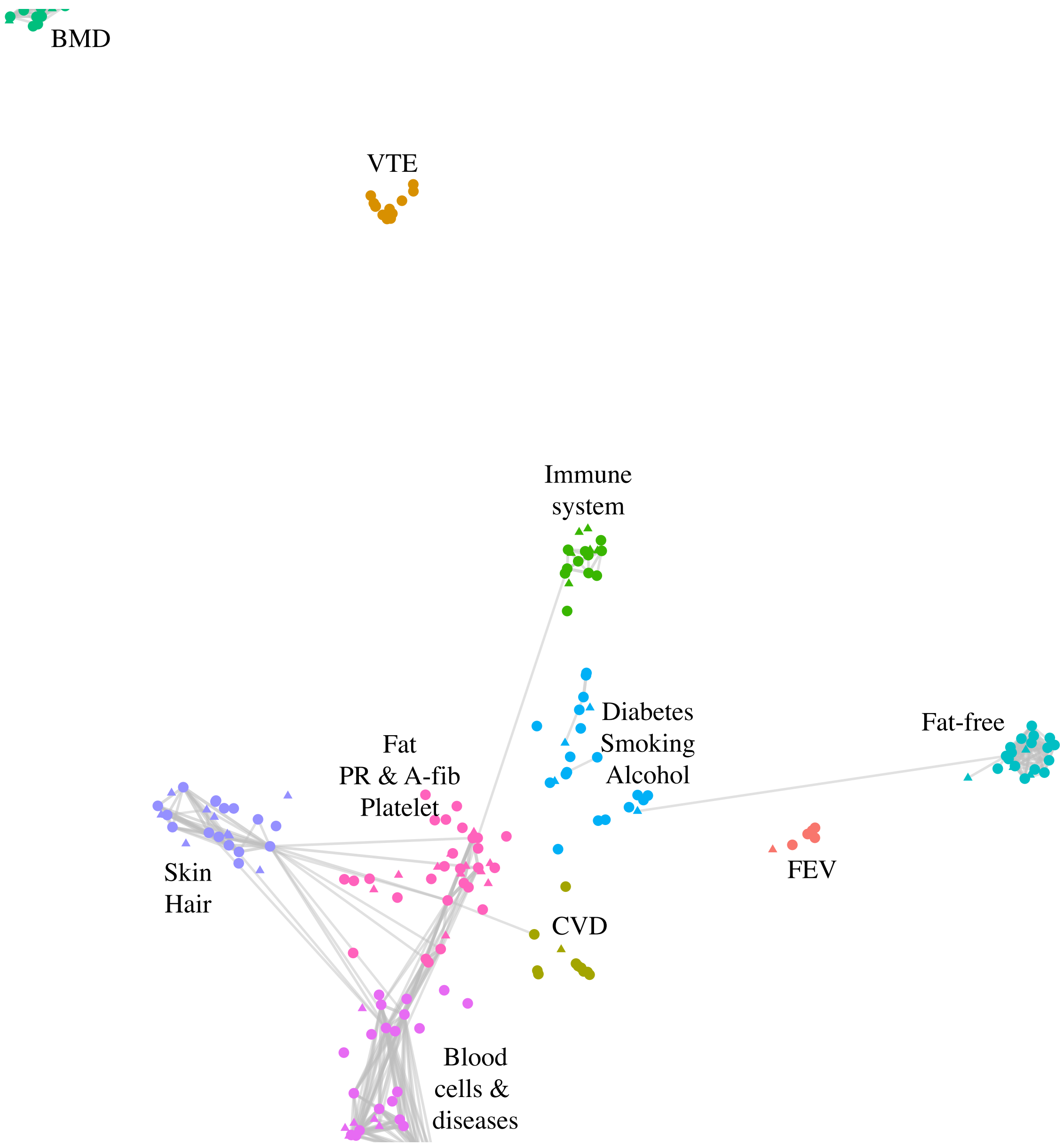

Figure 10: *Visualization of gene-trait clusters based on the UK biobank data using UMAP and kmeans. Circle points represent traits and triangle points represent candidate genes of signal index SNPs. A gene and a trait is connected by a grey line if the associated marginal association estimate is top 100 significant. We extract the major components accounting for $>$ 80% each cluster, summarize the traits, and annotate the clusters by the summaries. Abbreviations: BMD: bone mineral density; VTE: venous thromboembolism; Lymphs: Lymphocyte; FEV: forced expiratory volume; CVD: cardiovascular disease; RBC: red blood cell.*

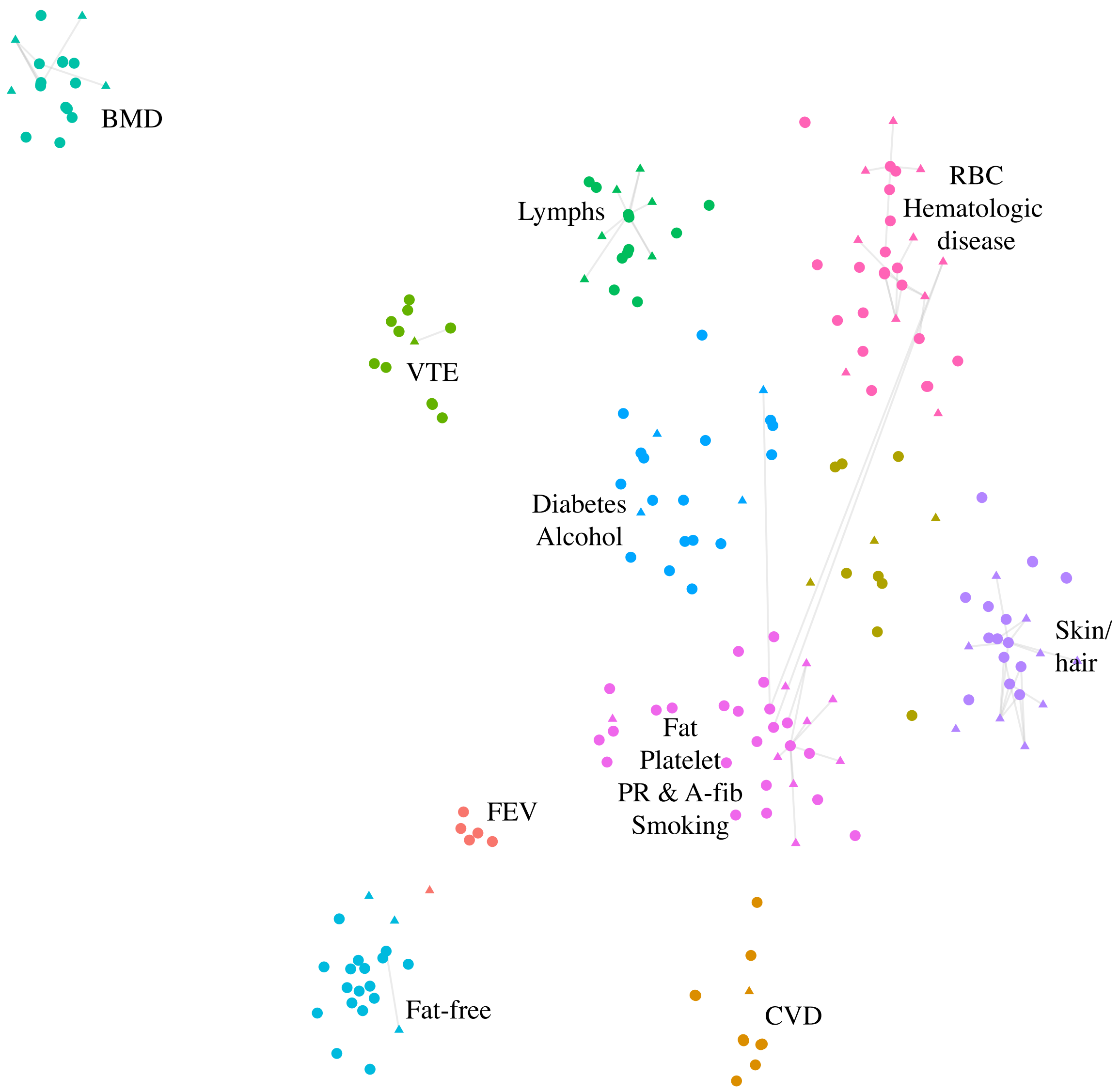

Figure 11: *Visualization of gene-trait clusters based on the UK biobank data using t-SNE and kmeans. More details can be found in the caption of Figure 10. We name the clusters according to their major components. Abbreviations: BMD: bone mineral density; VTE: venous thromboembolism; Lymphs: Lymphocyte; FEV: forced expiratory volume; CVD: cardiovascular disease; RBC: red blood cell; HR: heart rate.*

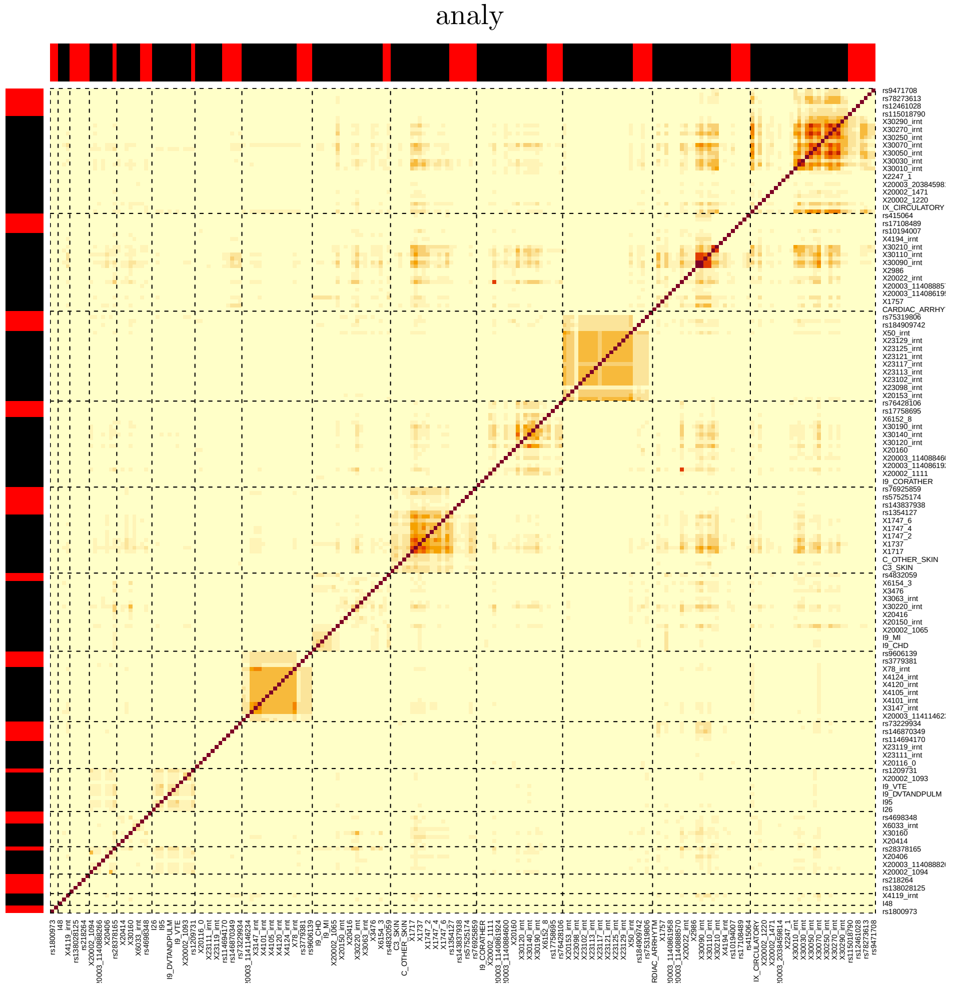

Figure 12: *Co-appearance heatmap based on the spectral clustering results on the UK biobank data. The dashed lines separate different clusters. The horizontal and vertical side bars use black to denote traits and red to denote SNPs.*

Since we do not have access to the true cluster memberships, we turn to pair-wise stability. In particular, for each pair $(a_i, a_j)$ ($a_i$, $a_j$ can be genes or traits), we compute the frequency over $m$ subsamples of the pair falling into the same cluster if $a_i$, $a_j$ both enter the analysis,

$$p_{ij} = \frac{\sum_{l=1}^{m} \mathbb{1}_{\{\exists\ C_k \in \mathcal{L}_l,\ a_i, a_j \in C_k\}}}{\sum_{i=1}^{m} \mathbb{1}_{\{a_i \in \mathcal{A}_l\}} \mathbb{1}_{\{a_j \in \mathcal{A}_l\}}}, \tag{22}$$

where in trial $l$ we denote the set of genes and selected traits by $\mathcal{A}_l$ and the gene-trait cluster list by $\mathcal{L}_l$.

Figure 13 plots histograms of pair-wise stability (22) over 100 trait subsampling based on the metabolomics data. We observe that co-appearance graph with bootstrap aggregation yields sharper spikes around zero and one, especially the latter. Close-to-one pairwise stability values indicate the associated pairs always fall into the same cluster and close-to-zero values imply the associated pairs always end up in different clusters. Thus, the "bathtub-shaped" curves of co-appearance graph with bootstrap aggregation suggest that a pair is always clustered or not clustered together, i.e., the clustering results are consistent across different sets of traits. Results based on the UK biobank data are similar and summarized in 14.

Figure 15 plots heatmaps of the pair-wise stability. Co-appearance graph with bootstrap aggregation produces heatmaps with more explicit structures while that of one-shot baseline is less clear. UMAP or t-SNE with k-means output less noisy heatmaps compared to spectral clustering. Results based on the UK biobank data are similar and summarized in 16.

*(a) One-shot baseline*

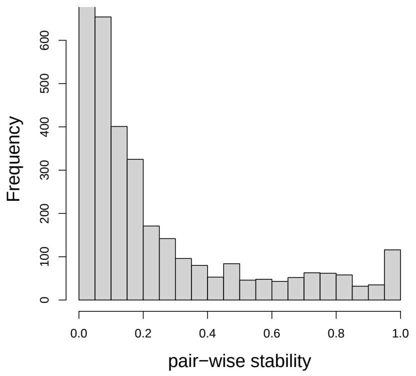

*(b) Co-apperance graph without bootstrapping*

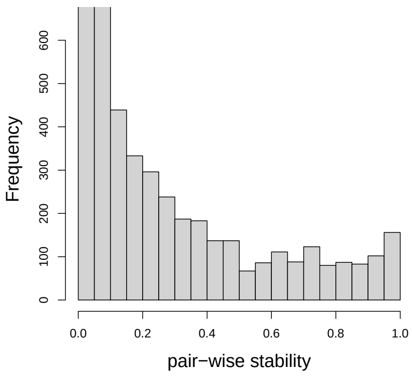

*(b.1) UMAP with k-means*

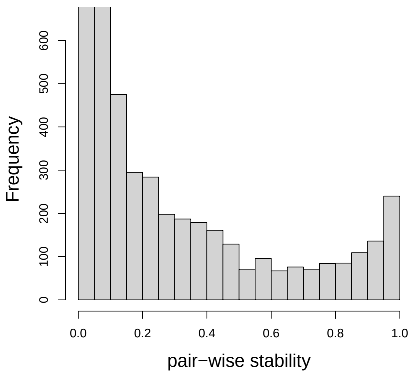

*(b.2) t-SNE with k-means*

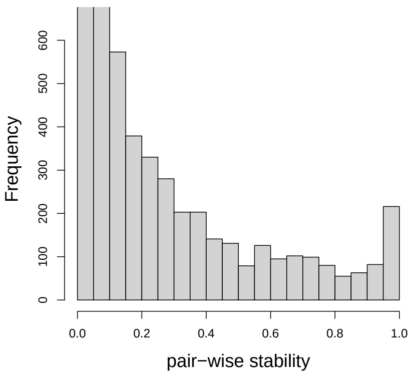

*(b.3) spectral clustering*

*(c) Co-apperance graph with bootstrapping*

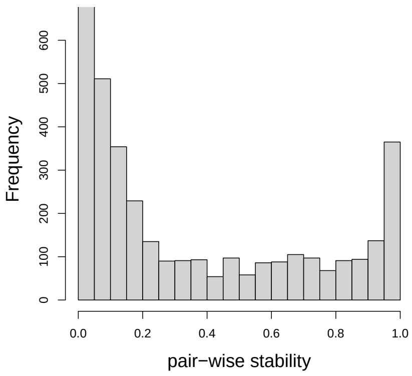

*(c.1) UMAP with k-means*

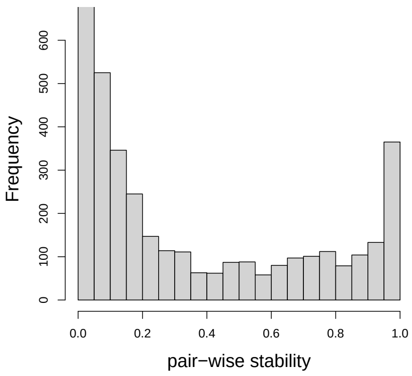

*(c.2) t-SNE with k-means*

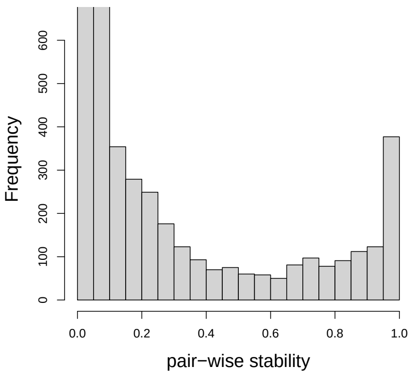

*(c.3) spectral clustering*

Figure 13: *Histograms of pair-wise stability* (22) *over* 100 *trait subsampling based on the metabolomics data. The first row represents one-shot baseline (Algorithm 1), the second row considers co-appearance graph without bootstrap aggregation and three clustering methods, and the third row does co-appearance graph clustering with bootstrap aggregation (Algorithm 2).*

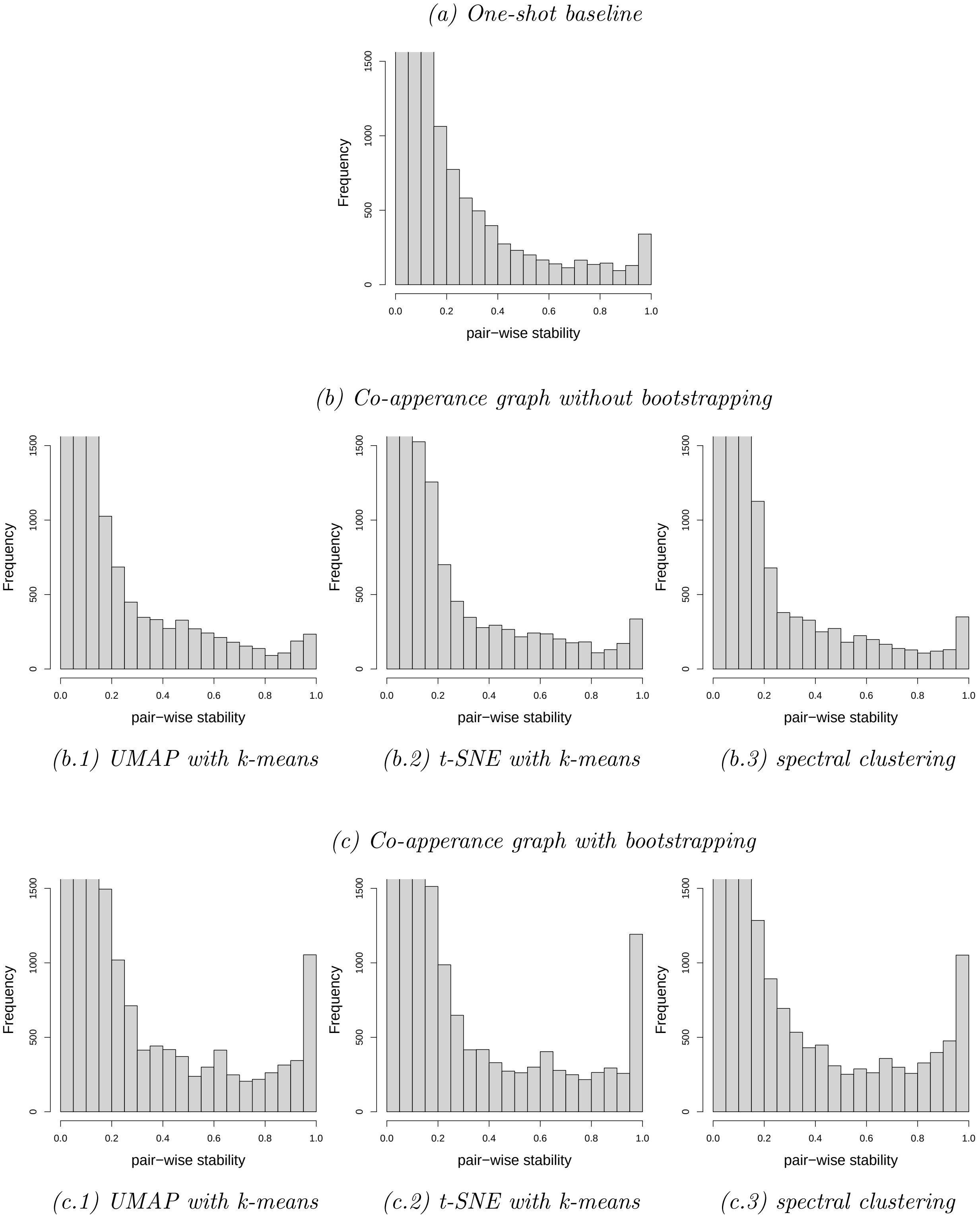

Figure 14: *Histograms of pair-wise stability* (22) *over* 100 *trait subsampling based on the UK biobank data. The first row represents one-shot baseline (Algorithm 1), the second row considers co-appearance graph without bootstrap aggregation and three clustering methods, and the third row does co-appearance graph clustering with bootstrap aggregation (Algorithm 2).*

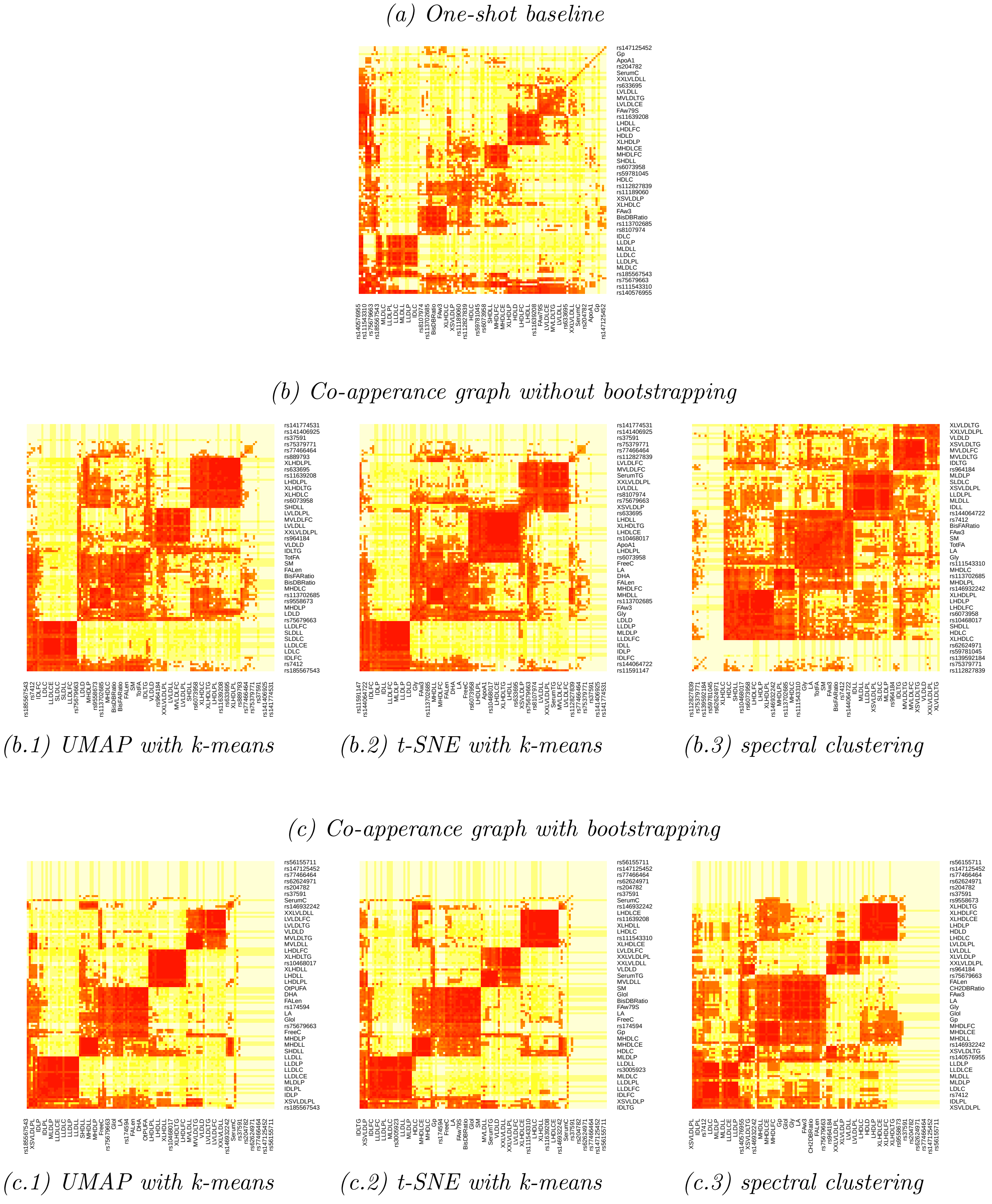

Figure 15: *Heatmaps of pair-wise stability (22) over 100 trait subsampling based on the metabolomics data. The first row represents one-shot baseline (Algorithm 1), the second row considers co-appearance graph without bootstrap aggregation and three clustering methods, and the third row does co-appearance graph clustering with bootstrap aggregation (Algorithm 2).)*

*(a) One-shot baseline*

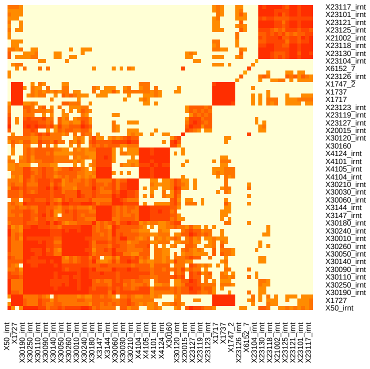

*(b) Co-apperance graph without bootstrapping*

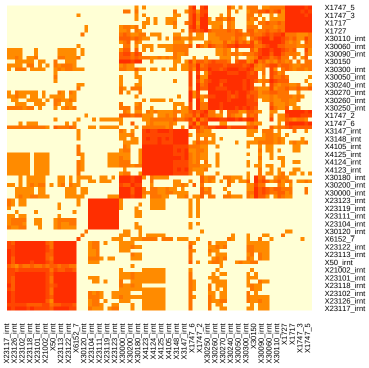

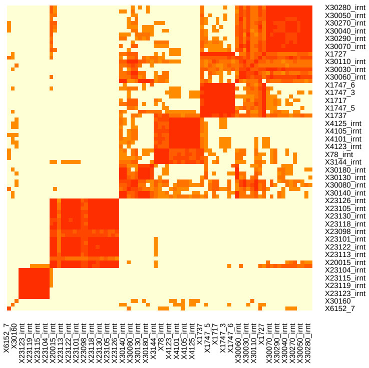

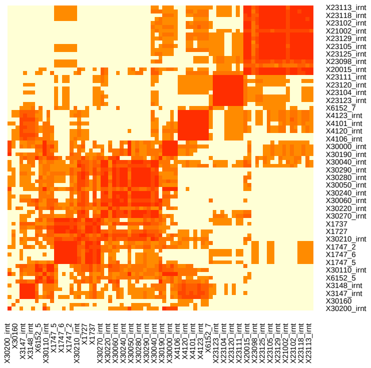

*(b.1) UMAP with k-means* *(b.2) t-SNE with k-means* *(b.3) spectral clustering*

*(c) Co-apperance graph with bootstrapping*

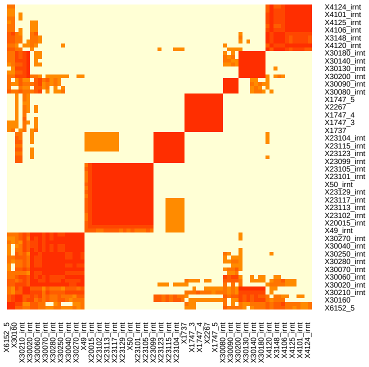

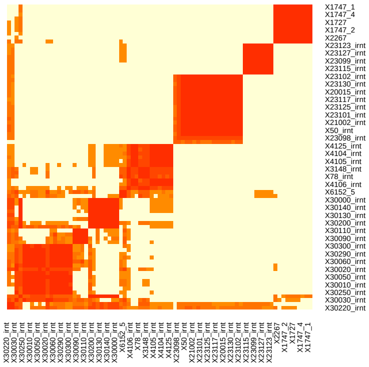

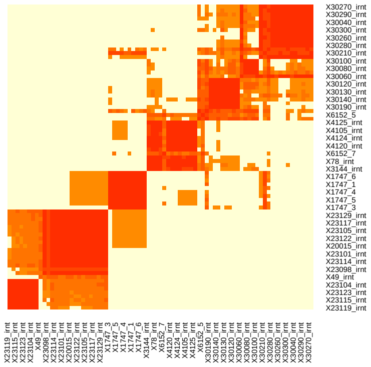

*(c.1) UMAP with k-means* *(c.2) t-SNE with k-means* *(c.3) spectral clustering*

Figure 16: *Heatmaps of pair-wise stability* (22) *over* 100 *trait subsampling based on the UK biobank data. The first row represents one-shot baseline (Algorithm 1), the second row considers co-appearance graph without bootstrap aggregation and three clustering methods, and the third row does co-appearance graph clustering with bootstrap aggregation (Algorithm 2).)*

| Cluster | Name | candidate gene | Organisms |
|---|---|---|---|
| L/M. VLDL | L.VLDL.C | | |
| | L.VLDL.CE | | |
| | L.VLDL.FC | | |
| | L.VLDL.PL | | |
| | L.VLDL.TG | | |
| | M.VLDL.C | | |
| | M.VLDL.CE | | |
| | M.VLDL.FC | | |
| | M.VLDL.L | | |
| | M.VLDL.P | | |
| | M.VLDL.PL | | |
| | M.VLDL.TG | | |
| | S.VLDL.TG | | |
| | VLDL.D | | |
| | XL.VLDL.TG | | |
| | Serum.TG | | |
| | FAw79S | | |
| | MUFA | | |
| | Tot.FA | | |
| | rs115849089 | AC100802.3 | |
| | rs1260326 | GCKR | Lymphoblastoid EUR exonlevel, Cells Transformed fibroblasts, Whole Blood |
| | rs140576955 | SNX18 | |
| | rs6976930 | BAZ1B | Thyroid, Skin Not Sun Exposed Suprapubic, Whole Blood |
| | rs8107974 | SUGP1 | Whole Blood |
| | rs964184 | ZNF259 | Lymphoblastoid EUR exonlevel |

Table 4: *Gene-trait clusters based on the metabolites data using UMAP and k-means (Part 1).*

| Cluster | Name | Candidate gene | Organisms |
|---|---|---|---|
| XL/L. HDL | L.HDL.C | | |
| | L.HDL.CE | | |
| | L.HDL.FC | | |
| | L.HDL.L | | |
| | L.HDL.P | | |
| | XL.HDL.C | | |
| | XL.HDL.CE | | |
| | XL.HDL.FC | | |
| | XL.HDL.L | | |
| | XL.HDL.P | | |
| | XL.HDL.PL | | |
| | XL.HDL.TG | | |
| | HDL.D | | |
| | ApoA1 | | |
| | rs10468017 | ALDH1A2 | |
| | rs111543310 | MYO1E | |
| | rs113083442 | RNF111 | |
| | rs11639208 | FAM63B | |
| | rs173539 | AC012181.1 | Cells Transformed fibroblasts, Whole Blood, Lung |
| | rs6073958 | PLTP | Lymphoblastoid EUR exonlevel, Muscle Skeletal, Adipose Visceral Omentum |
| | rs633695 | ALDH1A2 | |

*Gene-trait clusters based on the metabolites data using UMAP and k-means (Part 2).*

| Cluster | Name | Candidate gene | Organisms |
| --- | --- | --- | --- |
| LDL | LDL.C | | |
| | L.LDL.C | | |
| | L.LDL.CE | | |
| | L.LDL.FC | | |
| | L.LDL.L | | |
| | L.LDL.P | | |
| | L.LDL.PL | | |
| | M.LDL.C | | |
| | M.LDL.CE | | |
| | M.LDL.L | | |
| | M.LDL.P | | |
| | M.LDL.PL | | |
| | S.LDL.C | | |
| | S.LDL.L | | |
| | S.LDL.P | | |
| | rs11591147 | PCSK9 | |
| | rs12740374 | CELSR2 | Muscle Skeletal, Liver |
| | rs17248727 | LDLR | |
| | rs3005923 | AC1196-74.1 | |
| | rs7412 | APOE | Skin Not Sun Exposed Suprapubic |

*Gene-trait clusters based on the metabolites data using UMAP and k-means (Part 3).*

| Cluster | Name | Candidate gene | Organisms |
|---|---|---|---|
| Lipid measures | Bis.DB.Ratio | | |
| | Bis.FA.Ratio | | |
| | CH2.DB.Ratio | | |
| | CH2.in.FA | | |
| | DB.in.FA | | |
| | FALen | | |
| | FreeC | | |
| | Gln | | |
| | LA | | |
| | L.VLDL.P | | |
| | otPUFA | | |
| | PC | | |
| | SM | | |
| | Tot.PG | | |
| | ApoB | | |
| | rs174594 | FADS2 | Whole Blood, Lymphoblastoid EUR exonlevel |
| | rs56155711 | RP11-421P23.1 | |
| | rs75679663 | TM4SF5 | |

*Gene-trait clusters based on the metabolites data using UMAP and k-means (Part 4).*

| Cluster | Name | Candidate gene | Organisms |
|---|---|---|---|
| M/S. HDL | M.HDL.C | | |
| | M.HDL.CE | | |
| | M.HDL.FC | | |
| | M.HDL.P | | |
| | S.HDL.L | | |
| | S.HDL.P | | |
| | HDL.C | | |
| | LDL.D | | |
| | XL.VLDL.P | | |
| | Glol | | |
| | Gly | | |
| | Gp | | |
| | rs144064722 | ADAMTS3 | |

*Gene-trait clusters based on the metabolites data using UMAP and k-means (Part 5).*

| Cluster | Name | Candidate gene | Organisms |
|---|---|---|---|
| S/XS. VLDL | S.HDL.TG | | |
| | S.VLDL.FC | | |
| | S.VLDL.L | | |
| | S.VLDL.P | | |
| | S.VLDL.PL | | |
| | XS.VLDL.P | | |
| | XS.VLDL.TG | | |
| | IDL.TG | | |
| | FAw3 | | |
| | FAw6 | | |

*Gene-trait clusters based on the metabolites data using UMAP and k-means (Part 6).*

| Cluster | Name | Candidate gene | Organisms |
|---|---|---|---|
| IDL & XS. VLDL | IDL.C | | |
| | IDL.FC | | |
| | IDL.L | | |
| | IDL.P | | |
| | IDL.PL | | |
| | XS.VLDL.L | | |
| | XS.VLDL.PL | | |
| | Est.C | | |
| | rs185567543 | MTHFD2L | |

*Gene-trait clusters based on the metabolites data using UMAP and k-means (Part 7).*

| Cluster | Name | Candidate gene | Organisms |
|---|---|---|---|
| L/M. VLDL | L.VLDL.C | | |
| | L.VLDL.CE | | |
| | L.VLDL.FC | | |
| | L.VLDL.P | | |
| | L.VLDL.PL | | |
| | M.VLDL.C | | |
| | M.VLDL.CE | | |
| | M.VLDL.FC | | |
| | M.VLDL.L | | |
| | M.VLDL.P | | |
| | M.VLDL.PL | | |
| | M.VLDL.TG | | |
| | S.VLDL.TG | | |
| | VLDL.D | | |
| | FAw79S | | |
| | MUFA | | |
| | Serum.TG | | |
| | Tot.FA | | |
| | rs115849089 | AC1008-02.3 | |
| | rs140576955 | SNX18 | |
| | rs8107974 | SUGP1 | Whole Blood |
| | rs964184 | ZNF259 | Lymphoblastoid EUR exonlevel |

Table 5: *Gene-trait clusters based on the metabolites data using t-SNE and k-means (Part 1).*

| Cluster | Name | Candidate gene | Organisms |
|---|---|---|---|
| XL/L. HDL | L.HDL.C | | |
| | L.HDL.CE | | |
| | L.HDL.FC | | |
| | L.HDL.L | | |
| | L.HDL.P | | |
| | XL.HDL.C | | |
| | XL.HDL.CE | | |
| | XL.HDL.FC | | |
| | XL.HDL.L | | |
| | XL.HDL.P | | |
| | XL.HDL.PL | | |
| | XL.HDL.TG | | |
| | HDL.D | | |
| | ApoA1 | | |
| | rs10468017 | ALDH1A2 | |
| | rs111543310 | MYO1E | |
| | rs113083442 | RNF111 | |
| | rs11639208 | FAM63B | |
| | rs173539 | AC012181.1 | Lung, Cells Transformed fibroblasts, Whole Blood |
| | rs6073958 | PLTP | Lymphoblastoid EUR exonlevel, Adipose Visceral Omentum, Muscle Skeletal |
| | rs633695 | ALDH1A2 | |

*Gene-trait clusters based on the metabolites data using t-SNE and k-means (Part 2).*

| Cluster | Name | Candidate gene | Organisms |
|---|---|---|---|
| LDL | LDL.C | | |
| | L.LDL.C | | |
| | L.LDL.CE | | |
| | L.LDL.FC | | |
| | L.LDL.L | | |
| | L.LDL.P | | |
| | L.LDL.PL | | |
| | M.LDL.C | | |
| | M.LDL.CE | | |
| | M.LDL.L | | |
| | M.LDL.P | | |
| | M.LDL.PL | | |
| | S.LDL.C | | |
| | S.LDL.L | | |
| | S.LDL.P | | |
| | IDL.C | | |
| | IDL.FC | | |
| | IDL.L | | |
| | Est.C | | |
| | rs11591147 | PCSK9 | |
| | rs12740374 | CELSR2 | Muscle Skeletal, Liver |
| | rs17248727 | LDLR | |
| | rs3005923 | AC119674.1 | |
| | rs7412 | APOE | Skin Not Sun Exposed Suprapubic |

*Gene-trait clusters based on the metabolites data using t-SNE and k-means (Part 3).*

| Cluster | Name | Candidate gene | Organisms |
|---|---|---|---|
| Lipid measures | Bis.DB.Ratio | | |
| | Bis.FA.Ratio | | |
| | CH2.DB.Ratio | | |
| | CH2.in.FA | | |
| | DB.in.FA | | |
| | FALen | | |
| | FreeC | | |
| | Gln | | |
| | LA | | |
| | otPUFA | | |
| | PC | | |
| | SM | | |
| | M.HDL.CE | | |
| | rs174594 | FADS2 | Whole Blood, Lymphoblastoid EUR exonlevel |
| | rs56155711 | RP11-421P23.1 | |
| | rs75679663 | TM4SF5 | |

*Gene-trait clusters based on the metabolites data using t-SNE and k-means (Part 4).*

| Cluster | Name | Candidate gene | Organisms |
|---|---|---|---|
| M/S. HDL | M.HDL.C | | |
| | M.HDL.FC | | |
| | S.HDL.L | | |
| | S.HDL.P | | |
| | XL.VLDL.P | | |
| | LDL.D | | |
| | HDL.C | | |
| | Glol | | |
| | Gly | | |
| | Gp | | |
| | rs144064722 | ADAMTS3 | |
| | rs185567543 | MTHFD2L | |

*Gene-trait clusters based on the metabolites data using t-SNE and k-means (Part 5).*

| Cluster | Name | Candidate gene | Organisms |
|---|---|---|---|
| S/XS. VLDL | S.HDL.TG | | |
| | S.VLDL.FC | | |
| | S.VLDL.L | | |
| | S.VLDL.P | | |
| | S.VLDL.PL | | |
| | XS.VLDL.L | | |
| | XS.VLDL.P | | |
| | XS.VLDL.PL | | |
| | XS.VLDL.TG | | |
| | IDL.TG | | |
| | FAw3 | | |
| | FAw6 | | |

*Gene-trait clusters based on the metabolites data using t-SNE and k-means (Part 6).*

| Cluster | Name | Candidate gene | Organisms |
|---|---|---|---|
| IDL | IDL.P | | |
| | IDL.PL | | |
| | L.VLDL.TG | | |
| | M.HDL.P | | |
| | XL.VLDL.TG | | |
| | ApoB | | |
| | Tot.PG | | |
| | rs1260326 | GCKR | Lymphoblastoid EUR exonlevel, Cells Transformed fibroblasts, Whole Blood |
| | rs6976930 | BAZ1B | Thyroid, Skin Not Sun Exposed Suprapubic, Whole Blood |

*Gene-trait clusters based on the metabolites data using t-SNE and k-means (Part 7).*

| Cluster | Name | Candidate gene | Organisms |
|---|---|---|---|
| L/M. VLDL | L.VLDL.C | | |
| | L.VLDL.CE | | |
| | L.VLDL.FC | | |
| | L.VLDL.L | | |
| | L.VLDL.PL | | |
| | L.VLDL.TG | | |
| | M.VLDL.C | | |
| | M.VLDL.CE | | |
| | M.VLDL.FC | | |
| | M.VLDL.L | | |
| | M.VLDL.P | | |
| | M.VLDL.PL | | |
| | M.VLDL.TG | | |
| | S.VLDL.TG | | |
| | XL.VLDL.L | | |
| | XL.VLDL.TG | | |
| | VLDL.D | | |
| | FAw79S | | |
| | MUFA | | |
| | Serum.TG | | |
| | Tot.FA | | |
| | rs115849089 | AC1008-02.3 | |
| | rs140576955 | SNX18 | |
| | rs8107974 | SUGP1 | Whole Blood |
| | rs964184 | ZNF259 | Lymphoblastoid EUR exonlevel |

Table 6: *Gene-trait clusters based on the metabolites data using spectral clustering (Part 1).*

| Cluster | Name | Candidate gene | Organisms |
|---|---|---|---|
| XL/L. HDL | L.HDL.C | | |
| | L.HDL.CE | | |
| | L.HDL.FC | | |
| | L.HDL.L | | |
| | L.HDL.P | | |
| | L.HDL.PL | | |
| | XL.HDL.C | | |
| | XL.HDL.FC | | |
| | XL.HDL.L | | |
| | XL.HDL.P | | |
| | XL.HDL.PL | | |
| | HDL.D | | |
| | ApoA1 | | |
| | PC | | |
| | Tot.PG | | |
| | rs10468017 | ALDH1A2 | |
| | rs11639208 | FAM63B | |
| | rs633695 | ALDH1A2 | |

*Gene-trait clusters based on the metabolites data using t-SNE and k-means (Part 2).*

| Cluster | Name | Candidate gene | Organisms |
|---|---|---|---|
| LDL & IDL | LDL.C | | |
| | L.LDL.C | | |
| | L.LDL.CE | | |
| | L.LDL.FC | | |
| | L.LDL.L | | |
| | L.LDL.P | | |
| | L.LDL.PL | | |
| | M.LDL.C | | |
| | M.LDL.CE | | |
| | M.LDL.L | | |
| | M.LDL.P | | |
| | M.LDL.PL | | |
| | S.LDL.C | | |
| | S.LDL.L | | |
| | S.LDL.P | | |
| | IDL.C | | |
| | IDL.FC | | |
| | IDL.L | | |
| | IDL.PL | | |
| | ApoB | | |
| | Serum.C | | |
| | rs3005923 | AC119674.1 | |
| | rs7412 | APOE | Skin Not Sun Exposed Suprapubic |

*Gene-trait clusters based on the metabolites data using t-SNE and k-means (Part 3).*

| Cluster | Name | Candidate gene | Organisms |
|---|---|---|---|
| Lipid measures | Bis.DB.Ratio | | |
| | Bis.FA.Ratio | | |
| | CH2.DB.Ratio | | |
| | CH2.in.FA | | |
| | DB.in.FA | | |
| | DHA | | |
| | Est.C | | |
| | FALen | | |
| | FAw3 | | |
| | FAw6 | | |
| | FreeC | | |
| | LA | | |
| | otPUFA | | |
| | SM | | |
| | L.VLDL.P | | |
| | XL.VLDL.P | | |
| | rs174594 | FADS2 | Whole Blood, Lymphoblastoid EUR exonlevel |
| | rs75679663 | TM4SF5 | |

*Gene-trait clusters based on the metabolites data using t-SNE and k-means (Part 4).*

| Cluster | Name | Candidate gene | Organisms |
|---|---|---|---|
| M/S. HDL | M.HDL.C | | |
| | M.HDL.CE | | |
| | M.HDL.FC | | |
| | M.HDL.L | | |
| | M.HDL.P | | |
| | M.HDL.PL | | |
| | S.HDL.L | | |
| | S.HDL.P | | |
| | XL.HDL.CE | | |
| | XL.HDL.TG | | |
| | HDL.C | | |
| | LDL.D | | |
| | Gln | | |
| | Glol | | |
| | Gly | | |
| | Gp | | |
| | rs111543310 | MYO1E | |
| | rs144064722 | ADAMTS3 | |
| | rs173539 | AC012181.1 | Lung, Cells Transformed fibroblasts, Whole Blood |
| | rs185567543 | MTHFD2L | |

*Gene-trait clusters based on the metabolites data using t-SNE and k-means (Part 5).*

| Cluster | Name | Candidate gene | Organisms |
|---|---|---|---|
| S/XS. VLDL | S.VLDL.C | | |
| | S.VLDL.FC | | |
| | S.VLDL.L | | |
| | S.VLDL.P | | |
| | S.VLDL.PL | | |
| | XS.VLDL.L | | |
| | XS.VLDL.P | | |
| | XS.VLDL.PL | | |
| | XS.VLDL.TG | | |
| | XXL.VLDL.L | | |
| | IDL.P | | |
| | IDL.TG | | |
| | S.HDL.TG | | |
| | rs11591147 | PCSK9 | |

*Gene-trait clusters based on the metabolites data using t-SNE and k-means (Part 6).*

[GN18] Arjun K Gupta and Daya K Nagar. *Matrix variate distributions*, volume 104. CRC Press, 2018.

[HR02] Geoffrey Hinton and Sam T Roweis. Stochastic neighbor embedding. In *NIPS*, volume 15, pages 833–840. Citeseer, 2002.

[HW64] Alan E Hendrickson and Paul Owen White. Promax: A quick method for rotation to oblique simple structure. *British journal of statistical psychology*, 17(1):65–70, 1964.

[Kai58] Henry F Kaiser. The varimax criterion for analytic rotation in factor analysis. *Psychometrika*, 23(3):187–200, 1958.

[KDW+16] Johannes Kettunen, Ayşe Demirkan, Peter Würtz, Harmen HM Draisma, Toomas Haller, Rajesh Rawal, Anika Vaarhorst, Antti J Kangas, Leo-Pekka Lyytikäinen, Matti Pirinen, et al. Genome-wide study for circulating metabolites identifies 62 loci and reveals novel systemic effects of lpa. *Nature communications*, 7(1):1–9, 2016.

[LAE+21] Matthew S Lyon, Shea J Andrews, Ben Elsworth, Tom R Gaunt, Gibran Hemani, and Edoardo Marcora. The variant call format provides efficient and robust storage of gwas summary statistics. *Genome biology*, 22(1):1–10, 2021.

[MHM18] Leland McInnes, John Healy, and James Melville. Umap: Uniform manifold approximation and projection for dimension reduction. *arXiv preprint arXiv:1802.03426*, 2018.

[PNTB+07] Shaun Purcell, Benjamin Neale, Kathe Todd-Brown, Lori Thomas, Manuel AR Ferreira, David Bender, Julian Maller, Pamela Sklar, Paul IW De Bakker, Mark J Daly, et al. Plink: a tool set for whole-genome association and population-based linkage analyses. *American journal of human genetics*, 81(3):559–575, 2007.

[WK12] Lucas D Ward and Manolis Kellis. Haploreg: a resource for exploring chromatin states, conservation, and regulatory motif alterations within sets of genetically linked variants. *Nucleic acids research*, 40(D1):D930–D934, 2012.
